# Menin-Inhibition Sensitizes Acute Myeloid Leukemia to CLEC12A-Directed CAR Cell Therapy

**DOI:** 10.64898/2026.02.15.703376

**Authors:** Johanna Rausch, Philipp Wendel, Margarita Dzama-Karels, Marlene Steiner, Fenja Gierschek, Viktor Fetsch, Lea R. Knapp, Najla Abassi, Marie Kuhmann, Lilian Viehböck, Simon Weisemann, Nadezda Dolgikh, Jan Habermann, Catharina Lahrmann, Matthias Klein, Michael Delacher, Catherine Wölfel, Konstanze Döhner, Hartmut Döhner, Hakim Echchannaoui, Matthias Theobald, Daniel Sasca, Federico Marini, Robert Zeiser, Evelyn Ullrich, Michael W.M. Kühn

## Abstract

Menin inhibitors targeting the Menin-KMT2A chromatin complex have emerged as highly selective therapies for *KMT2A*-rearranged (*KMT2A*-r) and *NPM1*-mutated (*NPM1*^mut^) acute myeloid leukemia (AML), with recent regulatory approval and increasing interest in combination strategies. In contrast, CAR cell therapies have not yet been successfully established for AML. Here, we show that menin-inhibition primes *KMT2A*-r and *NPM1*^mut^ AML for CAR-based targeting by inducing robust and uniform expression of the myeloid antigen CLEC12A (CLL-1). Menin inhibitors did not impair T or NK cell viability, phenotype, or effector function. We engineered second-generation CLEC12A-directed CAR T cells that efficiently eliminated CLEC12A-positive AML. Across *in vitro* systems and xenograft models, the combination therapy consistently outperformed either monotherapy, resulting in profound disease control and significantly prolonged survival, with evidence of near-complete leukemia eradication *in vivo*. These findings support epigenetic priming with menin inhibitors to enhance CLEC12A-directed CAR cell-therapy in these AML subtypes.

**Significance:** Menin inhibitors, now approved for AML treatment, induce the immune target CLEC12A in *NPM1*^mut^ and *KMT2A*-r AML subtypes and sensitize AML cells to CLEC12A-directed CAR T cells without compromising immune function. As CLEC12A-CARs are already in clinical testing, this combination is immediately actionable for clinical investigation.

## Introduction

Epigenetic dysregulation is a hallmark of acute myeloid leukemia (AML) pathogenesis, and the majority of AML cases harbor disease-driving mutations in epigenetic regulators. Chromatin-based mechanisms frequently sustain leukemogenic transcriptional programs, thereby providing opportunities for targeted therapeutic intervention (reviewed in (1)).

Menin inhibitors are a recently developed class of small-molecule targeted agents that disrupt the interaction between menin and the chromatin regulator KMT2A. These agents were originally developed for *KMT2A*-rearranged (*KMT2A*-r) acute leukemias, in which chromosomal translocations generate oncogenic fusion proteins. KMT2A fusion proteins were initially shown to require menin for chromatin binding and for maintaining aberrant expression of self-renewal-associated *HOXA* and *MEIS1* transcription factor genes, which function as key leukemogenic drivers (2,3).

Using genome editing approaches, we previously identified the menin-KMT2A interaction as a critical dependency in *NPM1*-mutant (*NPM1*^mut^) AML (4). This most prevalent AML subtype, which is likewise driven by aberrant expression of *HOX* and *MEIS1* genes, is highly sensitive to menin-inhibition (menin-i). Early-generation menin inhibitors suppressed *HOX/MEIS1*-driven transcriptional programs and induced leukemic differentiation in preclinical models of *NPM1*^mut^ and *KMT2A*-r leukemias (2,4,5). Although *NPM1*^mut^ AML lacks KMT2A fusion proteins, subsequent studies demonstrated that mutant NPM1 aberrantly co-localizes with KMT2A to sustain menin-KMT2A-dependent *HOX* and *MEIS1* gene expression (6,7).

These preclinical findings motivated the development of clinical-grade menin inhibitors, which have since entered clinical evaluation (8–11). To date, seven distinct menin inhibitors entered clinical trials, both as monotherapies and in combination regimens, across multiple lines of AML treatment (12). Preliminary efficacy has been reported in phase I/II trials, with dramatic responses observed in heavily pretreated patients with relapsed or refractory (R/R) *NPM1*^mut^ or *KMT2A*-r acute leukemias treated with the menin inhibitors revumenib or ziftomenib (13–16). Based on these data, revumenib was approved by the U. S. Food and Drug Administration (FDA) for the treatment of R/R *NPM1*^mut^ and *KMT2A*-r acute leukemia, and ziftomenib for R/R *NPM1*^mut^ AML, representing the first targeted therapies approved for this most common AML subtype. Detailed molecular characterization of AML samples from patients treated in these studies confirmed that revumenib or ziftomenib treatment induces leukemic blast differentiation and silences HOX- and *MEIS1*-driven gene expression in responding patients (13,14). Although the genes directly regulated by the menin–KMT2A complex and suppressed by menin-i in *NPM1*^mut^ and *KMT2A*-r leukemias are well established through experimental models and clinical studies, it remains unclear which genes and proteins are activated by pharmacologic menin-i in these contexts. Moreover, it is unknown whether menin inhibitors induce specific cell-surface proteins that could be therapeutically exploited in combination strategies, including antibody-drug conjugates, bispecific T-cell engagers, or chimeric antigen receptor (CAR)-based therapies.

CAR immune effector cell therapy has significantly transformed cancer immunotherapy over the last decade, showing dramatic clinical efficacy against various lymphoid neoplasms (17–20), as reviewed in (21). CARs are genetically modified receptors that trigger a specific immune response against a target antigen, such as CD19 in lymphoid neoplasms (reviewed in (21)). Currently, seven different CAR T-cell products have been approved in the U.S. and Europe for the treatment of various lymphoid neoplasms, including acute lymphoblastic leukemia and diffuse large B-cell lymphoma (both targeting CD19) as well as multiple myeloma (targeting BCMA) (17–20) reviewed in(21)). However, developing CAR T cells for AML has proven challenging due to two main factors: 1) AML blasts often express target antigens that are also present on hematopoietic stem and progenitor cells; and 2) CAR products currently under clinical investigation for AML have shown limited efficacy, with early relapses likely resulting from variable expression of the target antigen on AML blasts and antigen escape (reviewed in (22)). At present, no CAR T-cell products for AML treatment are approved in the U.S. or Europe (22). Among potential target antigens, CD33, CD123, and C-type lectin domain family 12 member A (CLEC12A) are the most promising candidates. CAR T-cell products targeting CD33 and CD123 have undergone early-phase I clinical trials, but only limited preliminary efficacy has been reported for these approaches against AML (reviewed in (22). CLEC12A, also known as CLL-1, CD371, or MICL, is moderately expressed on AML cells and potentially on leukemic stem cells, while showing low or no expression on hematopoietic stem cells (HSCs), early progenitor cells (HPCs), and other body tissues (22–24). CLEC12A is a type II transmembrane receptor protein found primarily on monocytes, dendritic cells, and neutrophils, and it negatively regulates inflammatory pathways through its cytoplasmic domain (23,25). Initial clinical reports on CLEC12A-directed CAR therapies have reported promising complete remission rates in patients with relapsed or refractory AML. However, these responses are often followed by early relapses unless patients promptly undergo allogeneic stem cell transplantation (allo-SCT) (26–29).

In this paper, we demonstrate that menin-i induces uniform and robust upregulation of CLEC12A in *NPM1*^mut^ and *KMT2A*-r AML blasts, thereby priming them for next-generation CLEC12A-directed CAR T and NK cell-therapy, while preserving immune effector cell function.

## Results

### Menin inhibition induces remarkable CLEC12A expression in *NPM1*^mut^ and *KMT2A*-r AML

To explore whether menin inhibitors selectively induce cell-surface immune targets on AML cells, we revisited previously published RNA-sequencing (RNAseq) data from the *KMT2A*-r MOLM13 and MV411 cells and the *NPM1*^mut^ OCI-AML3 cells (5,30,31), upon treatment with different menin inhibitors vs. drug vehicle control (MI-503 (3µM), VTP-50469 (OCI-AML3 100nM; MV411 and MOLM13 50nM (30)) and KO-539 (150nM); Fig.1A). The analysis focused on upregulated genes that could serve as putative targets for a cellular therapy. Therefore, we first searched for genes that were uniformly induced by menin-i across all three AML cell lines and with all three menin inhibitors (using stringent cutoffs: log2-FC= 0.58, adj. p-value <0.05). Of the n=24 identified upregulated candidate genes (Fig.1A,B), we selected genes encoding surface receptors, narrowing the list to nine candidates (*TLR6, ST3GAL6, HLA-DQB1, NFAM1, PRLR, CLEC12A, MARCO, TNFSF138, TMEM229B*). Of these remaining nine genes, *CLEC12A* was the only cell-surface receptor gene reported to be commonly expressed on AML blasts, with minor or no expression on hematopoietic stem cells (HSCs), and no expression in any other organ tissue in the body according to *BloodSpot* and *the human protein atlas* databases (32,33), reducing the risk for severe hematologic or non-hematologic toxicities (22). Having identified *CLEC12A* as a menin inhibitor-inducible transcriptional target, we next assessed whether these transcriptional changes translate into the induction of the CLEC12A protein on the AML cell-surface using flow cytometry (FC). After seven-day VTP-50469 treatment (close analogue to revumenib), all *KMT2A*-r and *NPM1*^mut^ AML cells tested (MV411, MOLM13, OCI-AML3) revealed a significant increase in CLEC12A protein expression on the cell-surface (Fig.1C, Suppl.-Fig.1A+B, 3-8 fold MFI), while *KMT2A* and *NPM1* wild-type AML cell lines maintained their CLEC12A baseline expression (nearly 100% in HL60, nearly 0% in NB4). Similarly, six days of VTP-50469 treatment uniformly induced CLEC12A cell-surface expression in n=11 primary *NPM1*^mut^ AML patient samples (Fig.1D). On average, expression increased by 25.8%, with significant CLEC12A induction detected in all tested samples and the greatest induction in AML blasts with lower baseline expression (samples #1-7 with <60% CLEC12A baseline expression: induction of at least 35%).

**Figure 1:**
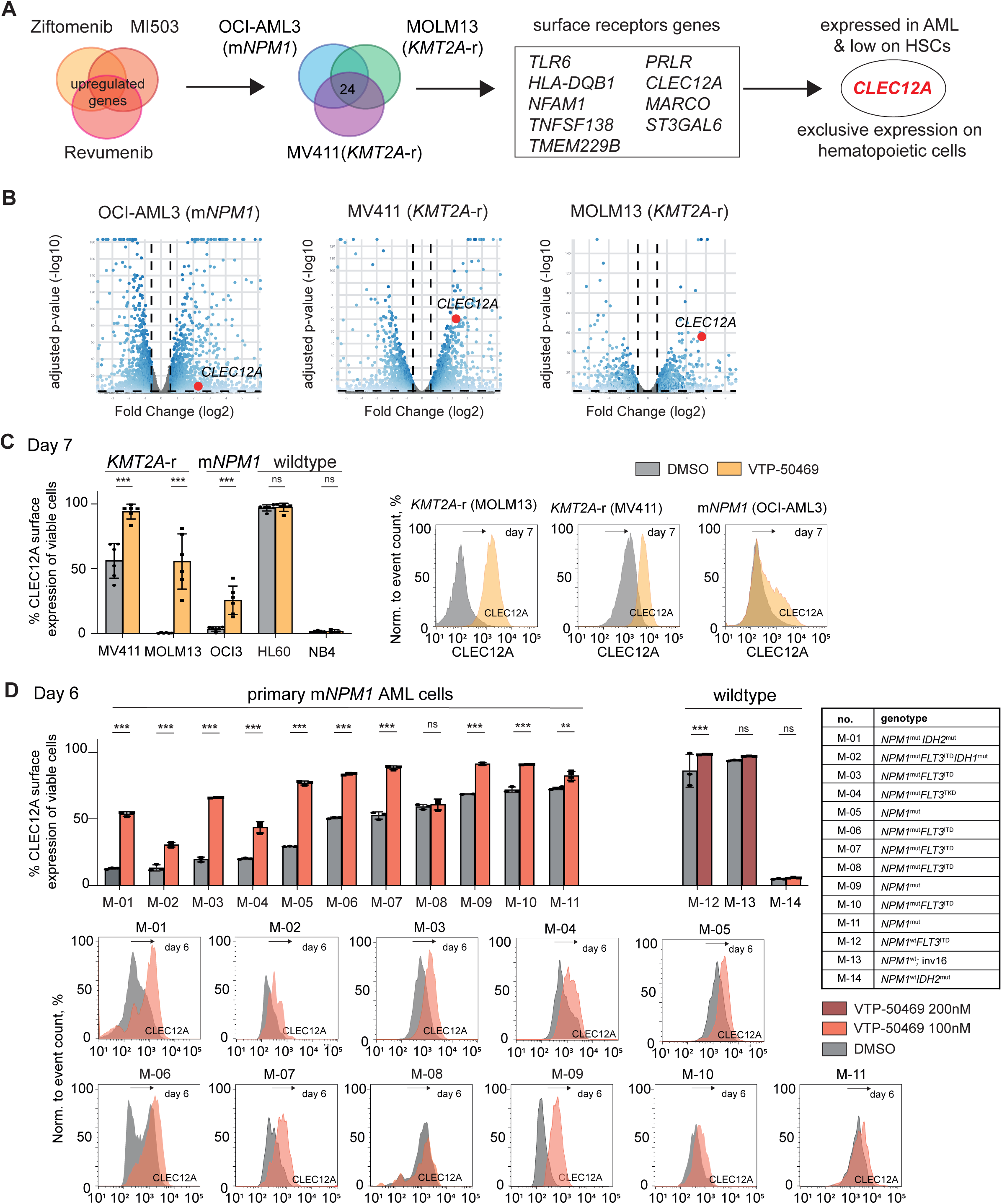
Menin inhibition uniformly induces the myeloid cell-surface antigen CLEC12A on *NPM1*^mut^ and KMT2A-r leukemic blasts. **(A) Identification of *CLEC12A* as a putative immune target induced by menin-inhibition.** RNAseq data of *KMT2A*-r MV411, MOLM13 and *NPM1*^mut^ OCI-AML3 cells treated with ziftomenib, VTP-50469 (close analogue revumenib) or MI-503 were selected for genes with significant induction (p_adj_ <0.05, log2-FC 0.58) upon treatment with all three menin inhibitors (Venn diagram on the left), and in all three cell lines (Venn diagram in the middle). Nine of these genes encode surface receptor proteins. CLEC12A was identified as the only target antigen with expression on leukemic blasts, low expression on hematopoietic stem cells and low expression in other organ tissues. **(B) *CLEC12A* gene expression is upregulated upon menin-inhibition in OCI-AML3, MV411 and MOLM13.** Volcano plots depicting the DESeq2-based differential expression analysis of RNAseq data comparing ziftomenib-treated (150nM) cells versus DMSO controls (OCI-AML3 and MOLM13 4-day treatment, MV411 3 days. Each symbol represents a gene, plotted by log2FC (ziftomenib versus DMSO) with p_adj_ <0.05. Marked is *CLEC12A*. **(C) CLEC12A surface expression is induced in *KMT2A*-r and *NPM1*^mut^ AML cell lines upon menin-inhibition.** *KMT2A*-r, *NPM1*^mut^ and wildtype cell lines were treated with 100nM VTP-50469 for 7 days. CLEC12A surface expression was determined by FC in viable (DAPI^low^) cells. Displayed is the mean of each experiment, all performed in technical triplicates. Significance by unpaired t-test, *** p<0.001. The histograms depict CLEC12A expression relative to the negative control, shown for one representative sample per sensitive cell line. **(D) CLEC12A surface expression is induced in *NPM1*^mut^ primary AML patient cells treated with menin-inhibition.** CLEC12A surface expression was determined by FC in primary AML patient cells with different genotypes (*NPM1*^mu*t*^ and *NPM1* wildtype, detailed information in the supplemental methods section) as indicated upon treatment with menin-i (VTP-50469 100nM or 200nM for 6 days) or DMSO. Significance by unpaired t test, ** p<0.01, *** p< 0.001. The histograms illustrate the induction of CLEC12A relative to the DMSO control after 6 days, shown for all *NPM1*^mut^ patient cells.

We then assessed CLEC12A expression in a model of non-genetic menin inhibitor resistance that we developed in our laboratory by increasing the drug concentration of MI-503 on single-cell clones of *NPM1*^mut^ OCI-AML3 cells (as described in the supplementary methods section). Of note, these cells, which do not harbor the well-described drug resistance mutations within the *MEN1* gene and exhibit a moderate increase in expression of monocytic differentiation markers, such as CD11b, maintain higher cell-surface CLEC12A expression levels than their normal counterparts (Suppl.-Fig.1C).

Next, we sought to explore what drives CLEC12A induction in AML cells upon menin-i. Therefore, we assessed the potential of menin and KMT2A chromatin binding to CLEC12A in Chromatin Immunoprecipitation followed by next-generation sequencing (ChIP-seq) data in *NPM1*^mut^ OCI-AML3 and previously published *KMT2A*-r MOLM13 cells (10), treated with DMSO or VTP-50469. As expected, we found no binding of menin or KMT2A to the CLEC12A locus in the DMSO or the VTP-50469 treated cells, confirming that CLEC12A is not a menin-KMT2A binding target (Suppl.-Fig.1D,E). As CLEC12A is normally expressed on neutrophils and monocytes, we speculated that the menin inhibitor-mediated differentiation may induce CLEC12A expression on AML blasts. To explore this hypothesis, we used all-trans retinoic acid (ATRA) as an alternate differentiation-inducing agent. CLEC12A expression was unstirred upon ATRA treatment in various AML cell lines, while cytomorphologic assessment confirmed the differentiating effect upon ATRA treatment, and CD11b was strongly induced (Suppl.-Fig.2A-C).

These findings demonstrate that menin-i uniformly induces CLEC12A surface expression in *NPM1*^mut^ and *KMT2A*-r leukemias but does not generally upregulate this protein upon treatment with other AML differentiating agents.

### Menin-Inhibition does not negatively influence immune effector cell function

Having established CLEC12A as an inducible antigen upon menin-i, we next characterized the general effects of pharmacological menin-i on immune effector cell functionality.

For *in vitro* menin inhibitor treatment assays, T cells were maintained in IL-2 and stimulated with agonistic anti-CD28/CD3 beads or irradiated OCI-AML3 cells, respectively, to mimic a leukemic environment for alloreactive stimulation. Natural Killer (NK) cells were stimulated with IL-15 (Fig.2A). To determine possible CLEC12A-directed CAR effector cell fratricide effects, we first assessed CLEC12A surface expression on CD3+ T and CD56+ NK cells upon 7, 14, and 21 days of menin inhibitor treatment. While CLEC12A expression on T cells was close to zero and did not significantly change with menin-i, a small proportion of cultured NK cells, derived from various donors, exhibited variable to low CLEC12A expression (on average: 16%, 33%, and 28%, day 7,14, 21) that did not increase with menin inhibitor treatment (Fig.2B).

**Figure 2:**
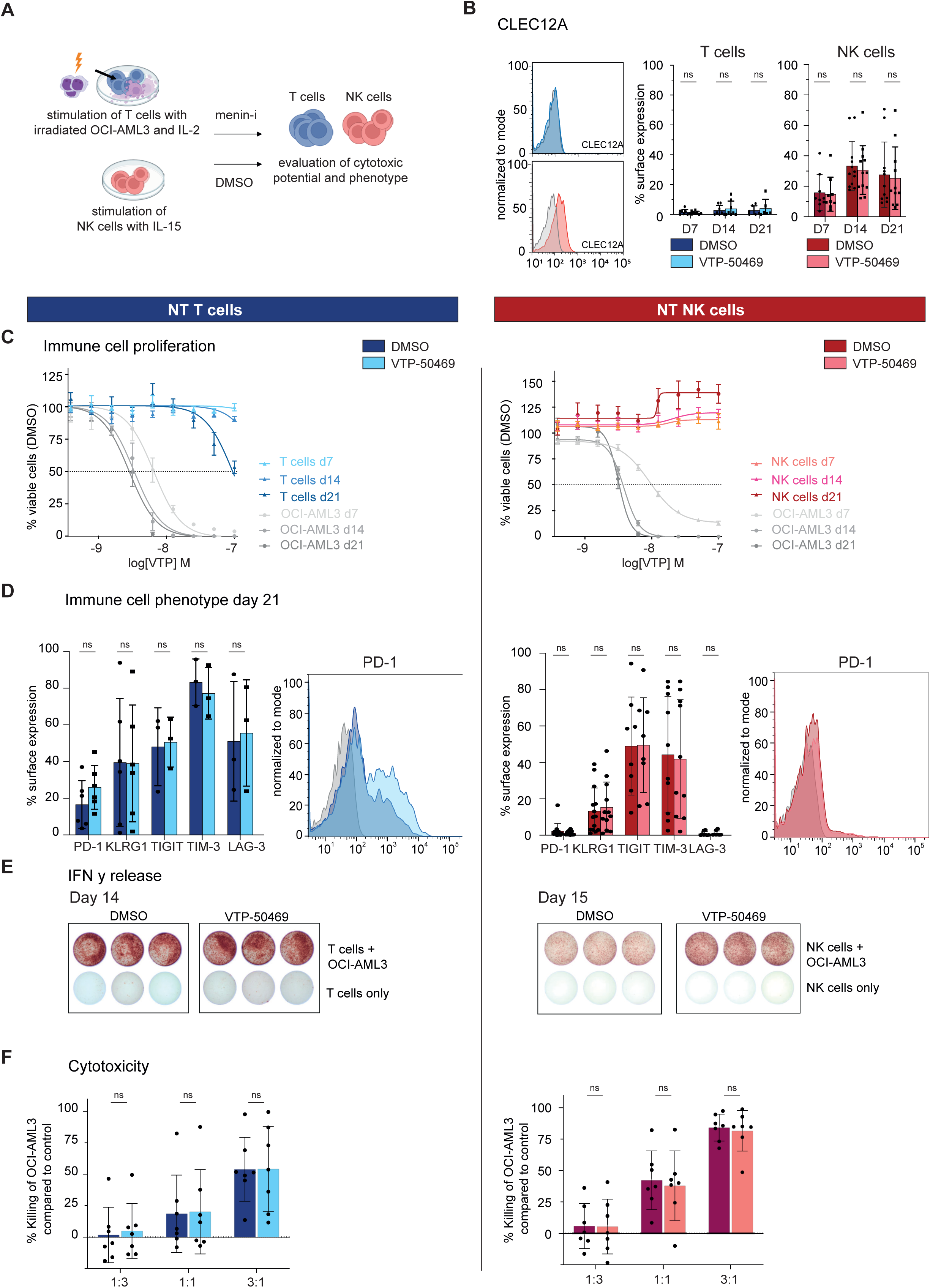
Menin-inhibition does not impair the functionality of non-transduced (NT) T or NK cells. **(A) Experimental design assessing the functionality of immune cells upon menin-inhibition.** NT NK cells were stimulated with IL-15, NT T cells with lethally irradiated OCI-AML3 and IL-2. Cells were treated with 100nM VTP-50469 starting day 0 and restimulated every 7 days. Figure made in BioRender.com. **(B) CLEC12A surface expression in NT T and NK cells is not affected by menin-inhibition.** FC-assessment of CLEC12A surface expression in viable T or NK cells after 7, 14, and 21 days of menin-i or vehicle control (DMSO). Displayed is the mean of each experiment, with statistical significance by an unpaired t-test. **(C) Minor effects of menin-inhibition on the proliferation of NT T and NK cells.** NK cells were stimulated with IL-15, T cells with agonistic anti-CD3/CD28 beads and IL-2. Treatment with different concentrations of VTP-50469 (100nM max, 1:2 dilutions or DMSO) for a total of 21 days. Displayed is the FC-readout of viable cells (DAPI^low^) on days 7, 14, and 21 compared to DMSO. OCI-AML3 cells were treated with the same concentrations of the menin inhibitor as a positive control. Summary of three experiments, each performed in technical triplicates, is displayed as the mean ± SEM for each concentration. **(D) Menin-inhibition has no relevant effect on the phenotype of NT T and NK cells.** Cells were treated as indicated in A. The surface expression of the immune checkpoints PD-1, KLRG-1, TIM-3, TIGIT and LAG-3 was determined on day 21 by FC. The histogram illustrates PD-1 expression in men-i compared to the DMSO control on day 21, shown exemplarily for one donor. **(E) IFN-γ response of NT T and NK cells is intact after menin inhibitor exposure.** IFN-γ release of native T and NK cells treated with VTP-50469 for 14 or 15 days was assessed by ELISpot assays after exposure to viable OCI-AML3 for 20h. **(F) Cytotoxic potential of NT T and NK cells is not affected by menin-inhibition.** Native T and NK cells were exposed to 100nM VTP-50469 for 7 days. Native T cells were simultaneously primed with lethally irradiated OCI-AML3 cells for 7 days for an alloreactive response. Immune cells were then co-cultured with viable OCI-AML3 cells at different E:T-ratios for 19h. Depicted is the percentage of killing relative to OCI-AML3 cells only (experimental design shown in Suppl.-Fig. 5E). Each symbol represents the mean of one experiment with a different biological immune cell donor; independent experiments were performed in technical triplicates. Significance by the unpaired t-test.

Next, we assessed whether the proliferative capacity of non-transduced (NT) T and NK cells was affected by menin inhibitor treatment. Over 21 days, immune effector cells were exposed to menin-i (VTP-50469), while proliferation was assessed every 7 days. T cell proliferation was not negatively affected after 7 and 14 days of continuous VTP-50469 exposure. Only after 21 days did we note a minor decrease in proliferation at very high concentrations of the drug compared to drug-vehicle (DMSO)-treated cells (Fig.2C, Suppl.-Fig.3A). NK cells modestly expanded over time (Fig. 2C).

We next explored the effects of menin inhibitors on immune cell phenotypes, IFN-γ response, and the cytotoxic efficacy of NT T and NK cells. The immunophenotype of T and NK cells was assessed using a panel of immune checkpoint markers (PD-1, KLRG-1, TIGIT, TIM-3, LAG-3) and did not reveal any changes upon 7, 14, and 21 days of menin-i (Fig.2D, Suppl.-Fig.3B) with PD-1 being the only exception showing a very mild, non-significant trend towards higher expression in the menin inhibitor-treated T cells (Fig.2D). Also, no significant changes were noted in menin inhibitor- vs. vehicle-treated T cells regarding the CD8/CD4-ratio or expression of CD62L, CD28, CD45RA, and CD57, which served as surrogates for the naïve and senescent-cell states (Suppl.-Fig.3B). In NK cells, the expression of the activation markers CD16, CD57, NKG2D and CD69 was not influenced by menin-i (Suppl.-Fig.4A).

To explore the IFN-γ response of T and NK cells upon menin-i, we pretreated the immune cells for 14 and 15 days with VTP-50469 before stimulation with viable OCI-AML3 cells and measured IFN-γ release using ELISpot assays 20 hours (h) later. While the secretion was not significantly changed in both cell types (Fig.2E, Suppl.-Fig.5A-D), there was a trend towards higher IFN-γ release in the NK cells upon menin-i (Fig.2E, Suppl.-Fig.5C,D).

Finally, we examined whether the cytotoxic potential of NT NK and T cells was affected by continuous exposure to menin-i. Therefore, T cells primed with lethally irradiated OCI-AML3 cells and NK cells cultured in the presence of IL-15 (Fig.2A) were assessed for their killing efficacy after seven days of menin-i. Therefore, the immune cells were co-cultured with luciferase (FLUC)-transduced OCI-AML3 cells at different Effector:Target (E:T)-ratios (Suppl.-Fig.5E), and killing was determined after 19h by comparing luminescence to that of the FLUC-OCI-AML3-only group (ctrl). No significant differences in T or NK cell killing efficacy were observed across all assessed E:T-ratios upon menin-i and vehicle control (Fig.2F). Overall, these analyses showed no significant adverse impact of menin-i on the functionality of T or NK cells.

### Immune effector cell subpopulations are proportionally maintained upon menin-inhibition

To determine whether menin-i induces proportional changes in the composition of immune effector cell subpopulations, we next performed single-cell RNA sequencing (scRNA-seq). CD3+ T cells and CD56+ NK cells were isolated from four independent donors and expanded for seven days in the presence of agonistic anti-CD3/CD28 beads and IL-2 for T cells, or IL-15 for NK cells, respectively. Cells were subsequently subjected to scRNA-seq following an additional four days of treatment with VTP-50469 (100nM) or DMSO control (Fig.3A). Transcriptional profiling resolved 17 distinct transcriptionally defined T cell subpopulations (Fig.3B, Suppl.-Fig.6A). Seven subpopulations corresponded to CD4+ T cells including four cytotoxic T-lymphocyte (CTL) clusters, two effector memory (EM) clusters, and one stem cell memory (SCM) cluster, while ten subpopulations corresponded to CD8+ T cells, comprising three central memory (CM) clusters, two EM clusters, one pre-effector cluster, one IL17RB-expressing effector cell cluster, and three effector T cell clusters exhibiting features of exhaustion (Fig.3B). All subpopulations were present under menin inhibitor- and vehicle-treated conditions without significant differences in relative abundance (Fig.3C). Consistently, differential gene expression analysis performed within each subpopulation did not identify any significantly altered genes between treatment and vehicle control (Suppl.-Fig.6B-D and Suppl.-Fig.7A-C)

**Figure 3:**
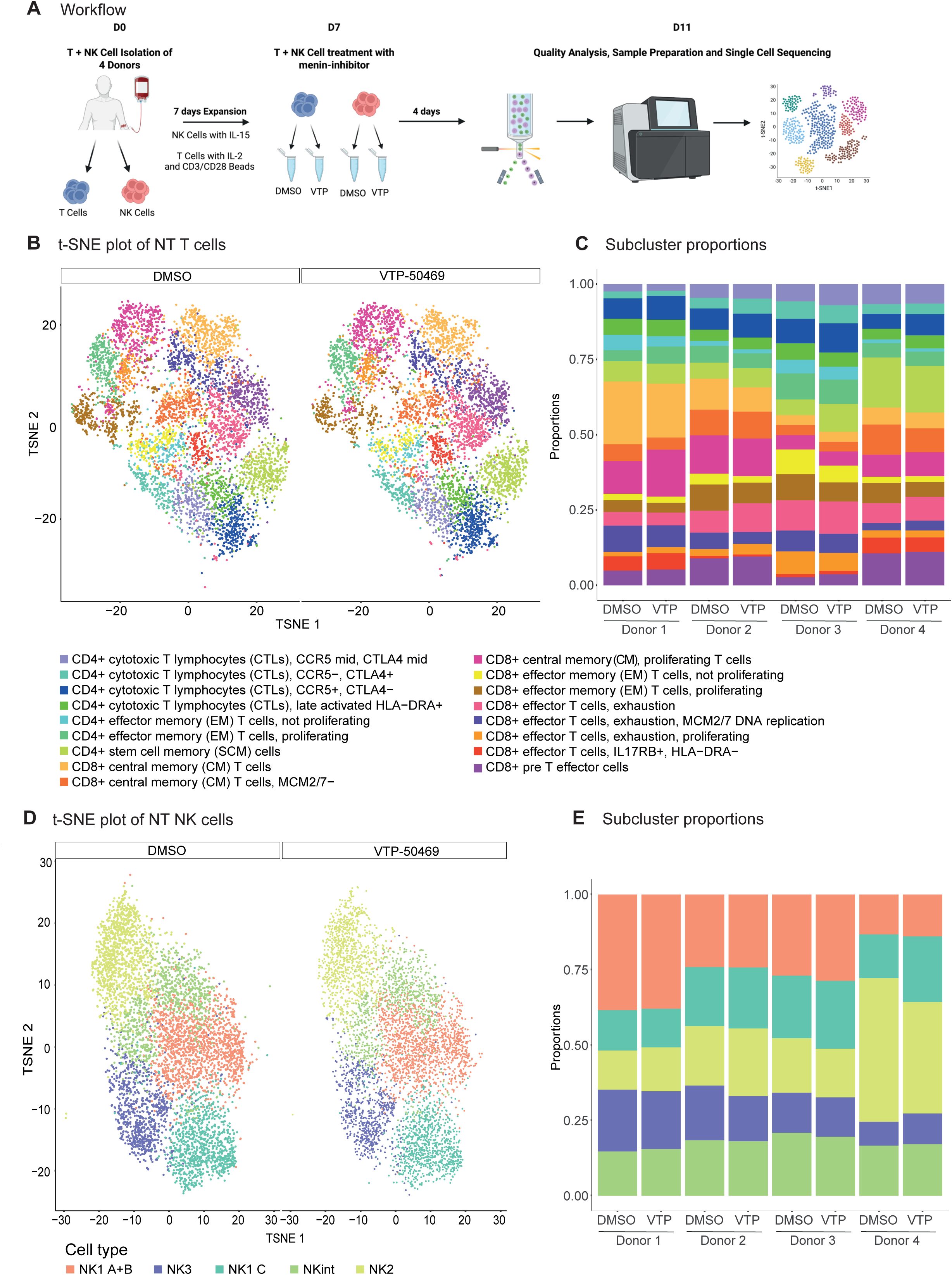
Single-cell sequencing reveals no changes within NT T or NK cell type proportions or differential gene expression upon menin-inhibition. **(A) Design of the single cell sequencing experiment.** NT NK or T cells from 4 different donors were isolated and expanded for 7 days with agonistic anti-CD3/CD28 beads and IL-2 for T cells or IL-15 for NK cells. Cells were exposed to 100nM VTP-50469 or DMSO for 4 days before processing for single-cell sequencing. Figure made in BioRender.com. **(B) t-SNE Plot of T cells.** 17 distinct T cell subpopulations were identified by clustering using the Louvain method. **(C) No differences in the proportions of NT T cell types.** Proportions of different T cell subtypes displayed for DMSO or menin-i for each donor. The proportion ratios between the two conditions across all four donors are close to 1 (0.82-1.36) with p-values for a difference >0.37. **(D) t-SNE Plot of NK cells.** 5 distinct NK cell clusters were identified using the Louvain algorithm. **(E) No differences in the proportions of NK cell types.** Proportions of different NK cell clusters displayed for DMSO or menin-i for each donor. The proportions ratios between the two conditions across all four donors are close to 1 (0.88-1.09) with p-values for a difference >0.52.

For NK cells, five major transcriptionally defined subpopulations (NK1A+B, NK1C, NK2, NKint, NK3) were identified (Fig.3D, Suppl.-Fig.8A), consistent with previously described NK cell states (34). None of these subpopulations exhibited significant changes in proportional representation following treatment (Fig.3E), and no significant treatment-associated differential gene expression was observed within individual NK cell subpopulations (Suppl.-Fig.8B-D, Suppl.-Fig.9). Collectively, these data indicate that menin-i does not induce substantial transcriptional remodeling in T or NK cell subpopulations.

### CLEC12A-engineered second-generation CAR T or NK cells effectively target CLEC12A-positive AML cells

After excluding negative effects of menin inhibitors on the functionality and transcriptionally defined composition of immune effector cell subpopulations, we next engineered a CLEC12A-directed CAR construct to evaluate its efficacy in combination with menin-i.

To this end, primary T and NK cells were genetically engineered via RD114TR- and VSV-G-pseudotyped lentiviral vectors encoding a second-generation, 4-1BB-based CLEC12A-CAR under the control of a human EF-1α promoter (Fig.4A,B) as described in detail in the methods and supplemental methods section. Stable and high CAR expression was confirmed by FC using the Myc-tag located in the hinge domain of the CAR and resulted in >88%±7% CAR+ cells for CAR T cells up to 17 days and 87%±4% for CAR NK cells 12 days post transduction (ptd) (Fig.4C). End-point analysis of immune cell subtypes on day 12-15ptd indicated no significant differences regarding the CD4+/CD8+ T cell populations and the CD16^dim^/CD16^bright^ NK cell population of CAR and NT immune cells (Fig.4D). In addition, the high purity of the analyzed immune cell populations was confirmed, with 99.5%±0.3% CD3+ for CAR T cells and 97%±2% CD56+ for CAR NK cells (Fig.4D).

**Figure 4:**
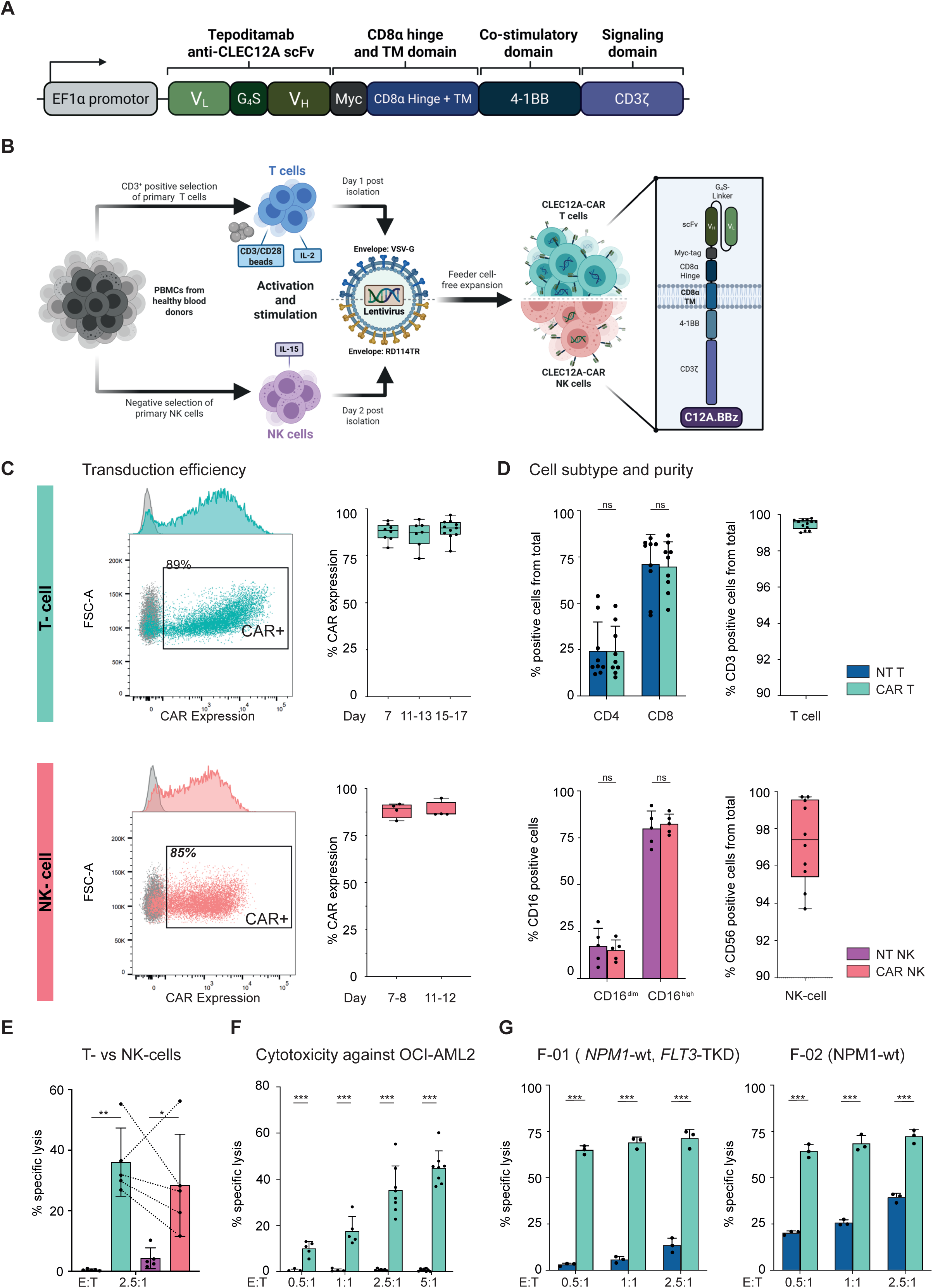
Enhanced functionality of CLEC12A-CAR immune cells against AML cells *in vitro*. **(A) CAR design.** Schematic representation of the developed second-generation CLEC12A-CAR. Figure made in BioRender.com. **(B) Production workflow for the generation of primary CAR T and CAR NK cells.** Primary immune cells were enriched from healthy donor PBMCs, activated with IL-2 and CD3/CD28 beads (T cells) or low-dose IL-15 (NK cells) and transduced using VSV-G- and RD114TR-pseudotyped lentiviral vectors at the indicated timepoints. Figure made in BioRender.com. **(C) CAR T and CAR NK cells demonstrate a high CLEC12A-CAR expression.** CLEC12A-CAR surface presentation detected *via* Myc-Tag 11-12 days post transduction (ptd) for one representative donor and monitored CAR expression during cultivation over up to 17 days (T cells, n=7-9) and 11 days (NK cells, n =4). The histogram on the left depicts the transduction efficacy, shown exemplarily for one donor. **(D) High purity of CAR T and NK cells with similar cell subtypes compared to their NT counterparts.** Phenotype of expanded CAR NK and CAR T cells was determined *via* FC based on CD16^dim^/CD16^high^ (NKs cell, n=5) and CD4+/CD8+ (T cells, n=9) subpopulations 12-15 days ptd. Purity of NT and CAR immune cells based on CD3+ (T cells, n=13) and CD56+/CD3- cells (NK cells, n=10) 12-15 days ptd. **(E) CAR T and NK cells exhibit high cytotoxic functionality.** Comparison of CAR immune cell functionality of matching donors using a 4h, FC-based cytotoxicity assay against OCI-AML2 cells at E:T-ratio 2.5:1 (n=5) on day 11-13ptd. Dotted lines indicate matching CAR immune cell donors. Significance by unpaired t-tests * p<0.05, ** p<0.01 **(F) CAR T cells show strong cytotoxicity against OCI-AML2 cells.** CLEC12A-CAR T cell functionality was investigated against OCI-AML2 following 4h co-culture at increasing E:T-ratios 0.5:1 to 5:1 (n=5-8). Significance by multiple unpaired t-tests *** p<0.001. **(G) CAR T cells effectively target primary AML patient cells.** Superior efficacy of CAR T cells compared to NT T cells against primary AML patient cells, following co-culture for 24 at E:T-ratio 0.5:1 to 2.5:1 (n=3). Significance by multiple unpaired t-tests *** p<0.001

The functionality of CLEC12A-CAR immune cells derived from matching donors was then compared in FC-based cytotoxicity assays against the *KMT2A*-r OCI-AML2 cells, which we found to have very high baseline CLEC12A expression. Co-cultivation of the CAR immune effector cells with OCI-AML2 for 4h at an E:T-ratio of 2.5:1 resulted in strong cytotoxicity. We observed a significant increase in specific tumor cell lysis, with mean values of 36% for CAR T cells and 28% for CAR NK cells, compared to NT T and NT NK cells, respectively (Fig.4E). Notably, comparisons of matched immune cell donors showed a slight trend toward increased functionality of CLEC12A-CAR T cells compared to CAR NK cells. CAR T cell functionality was further investigated in cytotoxicity assays against OCI-AML2 at E:T-ratios from 0.5:1 to 5:1, resulting in significantly enhanced tumor cell lysis at an E:T-ratio of 5:1, with up to 42%±4% lysis after 4h and complete lysis of 95%±2% within 24h (Fig.4F; Suppl.-Fig.10A). Subsequently, CLEC12A-directed CAR T cells were challenged with two primary AML patient samples (F-01 *NPM1*^WT^*FLT3*^TKD^, F-02 *NPM1*^WT^) that we selected based on very high CLEC12A baseline expression (86% and 89%) (Fig.4G, Suppl.-Fig.10B,C). These experiments demonstrated very high CAR T cell efficacy, with AML-specific lysis exceeding 60% after 24h of coculture at an E:T-ratio of 2.5:1 (Fig.4G). These data demonstrate that the here-engineered CLEC12A-directed CAR T cells are highly effective against AML blasts with high baseline CLEC12A expression and provide a strong basis for use in a combinatorial approach with menin-i.

### Menin-inhibition primes *NPM1*^mut^ and *KMT2A*-r AML for more efficient CLEC12A-directed CAR effector cell therapy

Our data are consistent with menin-i uniformly upregulating CLEC12A in *NPM1*^mut^ and *KMT2A*-r AML blasts. After successfully engineering highly functional CLEC12A-directed CAR T and CAR NK cells, we sought to investigate the effects of combining menin-i with these CLEC12A-directed CAR effector cells. First, we pretreated *FLUC*-transduced OCI-AML3 cells with VTP-50469 or vehicle control for four days (Fig.5A) and then confirmed CLEC12A induction in the VTP-50469-treated cells (Suppl.-Fig.11A). Next, equal numbers of DMSO- or menin inhibitor-pretreated FLUC-OCI-AML3 cells were replated, and then CAR effector cells were added at different E:T-ratios (Fig.5A). The killing efficacy of the CLEC12A-directed CAR effector cells against menin inhibitor- versus vehicle-treated FLUC-OCI-AML3 cells was quantified after 19h using a luminescence-based viability assay.

**Figure 5:**
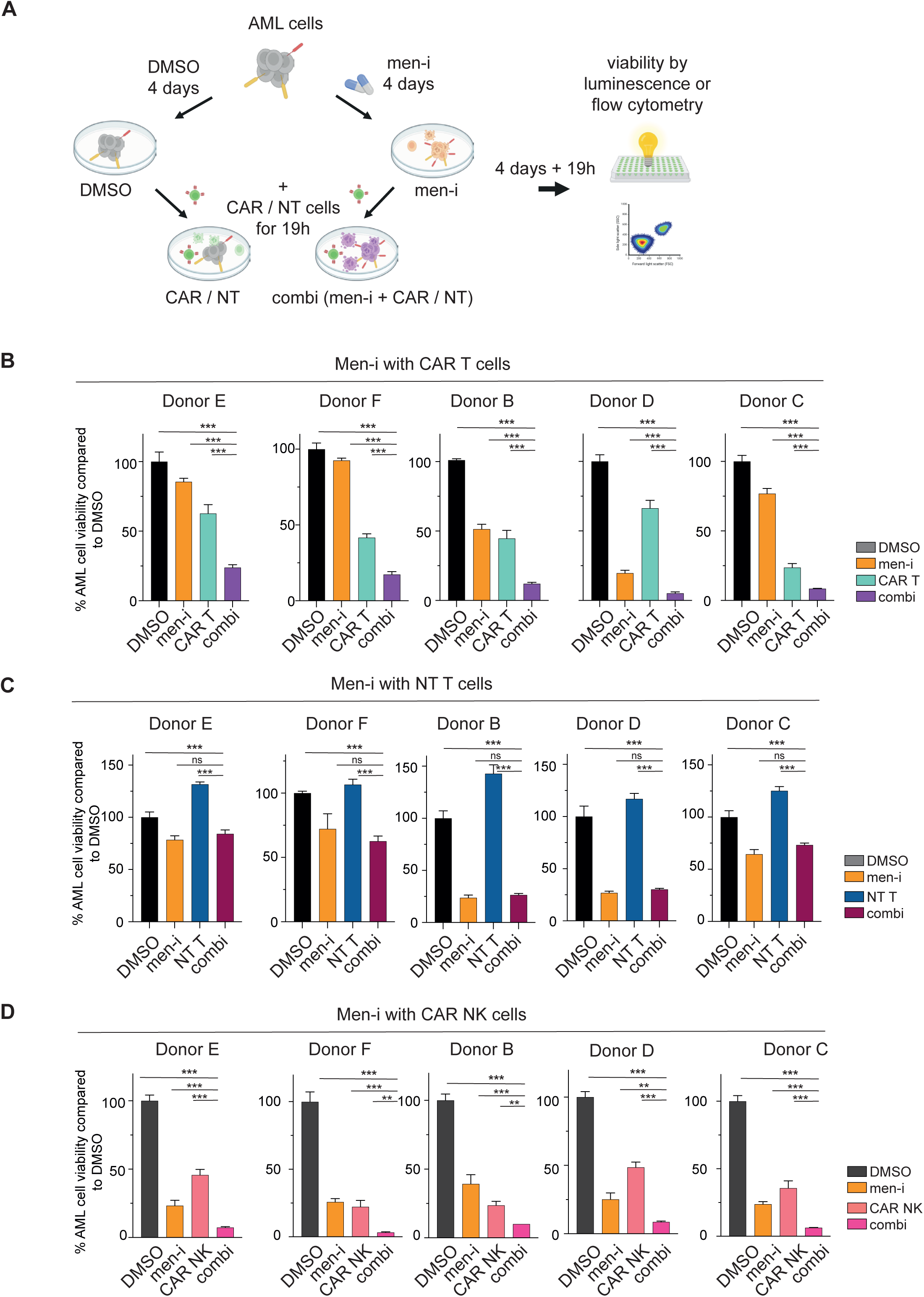
The combination of menin-inhibition and CLEC12A-directed CAR therapy outcompetes the anti-leukemic activity compared to single treatments *in vitro*. **(A) Design of the combination assay.** *FLUC*-OCI-AML3 were treated with 100nM VTP-50469 or DMSO for 4 days. On day 4, cells were counted, and the same number of DMSO- or menin inhibitor-treated viable AML cells were plated, either stained with CSFE or incubated with D-Luciferin in the media. NT or CAR cells were added at different E:T-ratios to the pretreated AML cells in the respective groups (CAR/NT and combi). Viability was determined after 19h relative to the DMSO group. Figure made in BioRender.com. **(B) Combination of menin-inhibition and CAR T-cell therapy results in superior anti-leukemic effects compared to single treatments.** *FLUC*-OCI-AML3 cells were exposed to 100nM VTP50469 or DMSO as previously described, and CAR T cells were added as effector cells. Displayed is the E:T-ratio of 1:1, both for the single treatment and the combination with menin-i. Five independent experiments with different biological donors, each experiment in technical triplicates. AML cell viability is displayed, with killing defined as 100% − viability. Significance by unpaired t-test ** p<0.01, *** p<0.001. **(C) NT T cells exhibit no specific lysis nor additive anti-leukemic effects in combination with menin-inhibition.** *FLUC*-OCI-AML3 cells were exposed to 100nM VTP50469 or DMSO as previously described, and NT T cells were added as effector cells. Displayed is the E:T-ratio of 1:1, both for the single treatment and the combination with menin-i. Five independent experiments with different biological donors, each experiment in technical triplicates. AML cell viability is displayed, with killing defined as 100% − viability. Significance by unpaired t-test *** p<0.001, not significant (ns). **(D) Combination of menin-inhibition and CAR NK-cell therapy exhibits superior anti-leukemic effects.** *FLUC*-OCI-AML3 cells were exposed to 100nM VTP50469 or DMSO as previously described, and CAR NK cells were added as effector cells. Displayed is the E:T-ratio of 1:1, both for the single treatment and the combination with menin-i. Five independent experiments with different biological donors, each experiment in technical triplicates. Significance by unpaired t-test * p<0.05, ** p<0.01, *** p<0.001.

Therefore, we added CLEC12A-directed CAR T cells to menin inhibitor-pretreated OCI-AML3 cells and observed significantly enhanced AML killing with the combination versus both single treatments across all individual donors and different E:T-ratios (Fig.5B, Suppl.-Fig.11B,C). No AML killing was observed for the NT T cells and the combination showed similar killing as menin-i alone (Fig.5C, Suppl.-Fig.11D,E). CLEC12A-CAR T cell specific lysis compared to the NT T cells was on average 87% vs. 45% for the CAR T vs. NT T cells (E:T-ratio 1:1), both in combination with menin-i (menin-i cell killing alone 35% in the CAR T cell assay (p=0.0035) and 47% in the NT T cell assay (statistically non-significant)) (Fig.5B+C, Suppl.-Fig.11B-E). Notably, the superiority of the combination of CAR T cells and menin-i was also observed after 6h of co-culture with OCI-AML3 target cells in higher E:T-ratios (Suppl.-Fig.11F,G).

To assess whether CAR T cell-specific AML killing increased with the menin inhibitor-induced CLEC12A expression, we compared CLEC12A-CAR T cell-mediated killing between menin inhibitor- and DMSO-pretreated AML cells. These analyses confirmed enhanced CAR T cell killing of menin inhibitor- versus DMSO-pretreated AML cells, more pronounced for the higher E:T-ratios (Suppl.-Fig.11C). Next, we confirmed the significantly superior AML killing of combined menin-i with CLEC12A-CAR T cell treatment against the *KMT2A*-r MOLM13 AML cells that exhibit an even more pronounced induction of CLEC12A expression upon menin-i from a low baseline level (Fig.1C, Suppl.Fig.1A,B). Again, the combination of CAR T cells with menin-i was superior to single treatments, although with substantial variability between different T cell donors (Suppl.-Fig.12,A-F).

The combination of CLEC12A-directed CAR NK cells with menin-i also substantially improved AML killing compared to single drug treatments across all donors (Fig.5D, Suppl.-Fig.13A-C). However, NT NK cells showed similar anti-leukemic efficacy as their CLEC12A-directed counterparts, possibly due to their strong ability to HLA-independent killing of OCI-AML3 (Suppl.-Fig.13D,E), as reported previously (35).

As NK cells show variable low-level expression of the immunotarget CLEC12A, it is interesting to note that CLEC12A self-expression on the transduced CAR NK cells vs. NT NK cells – both derived from the same donor - was substantially lower within the CAR NK cells, consistent with a potential fratricide-induced selection for CLEC12A non-expressing CAR NK cells (Suppl.-Fig.13F). As shown in Fig.2A CLEC12A expression was very low to absent on T cells and remained below 1% in the CLEC12A-CAR T cells (Suppl.-Fig.13G).

Next, we also assessed whether CAR T and CAR NK cells exhibited immunophenotypic changes-possibly associated with immune cell exhaustion- compared to their NT counterparts, but did not find significant changes in the expression of CD4, CD8, PD-1, or KLRG-1 on T cells or CD16, PD-1, or KLRG-1 on NK cells upon seven days of menin-i (Suppl.-Fig.13F, G).

Together, these data suggest that combinatorial menin-i with CLEC12A-directed CAR T cells has substantially enhanced therapeutic effects against *NPM1*^mut^ and *KMT2A*-r AML, likely conveyed by uniform menin inhibitor-induced CLEC12A target expression on leukemic blasts, resulting in greater CLEC12A-CAR directed killing.

### The combination of CLEC12A-directed CAR T cells and menin-inhibition has profound activity against primary AML cells and can eradicate disease in an aggressive human *KMT2A*-r AML xenograft model

Next, we assessed the combined CLEC12A-CAR plus menin-i treatment regimen in primary *NPM1*^mut^ AML cells. The primary AML samples were cultured for 6 days with 100nM VTP50469 in serum-free media as previously described (5) before being exposed to CLEC12A-engineered CAR T cells for 19h. Again, the anti-leukemic effects of the combination were superior to those of every single treatment, reaching statistical significance in three of the five samples (Fig.6A). Of note, all samples with significantly superior anti-leukemic activity demonstrated a low CLEC12A expression upon DMSO exposure and strong induction upon menin-i).

**Figure 6:**
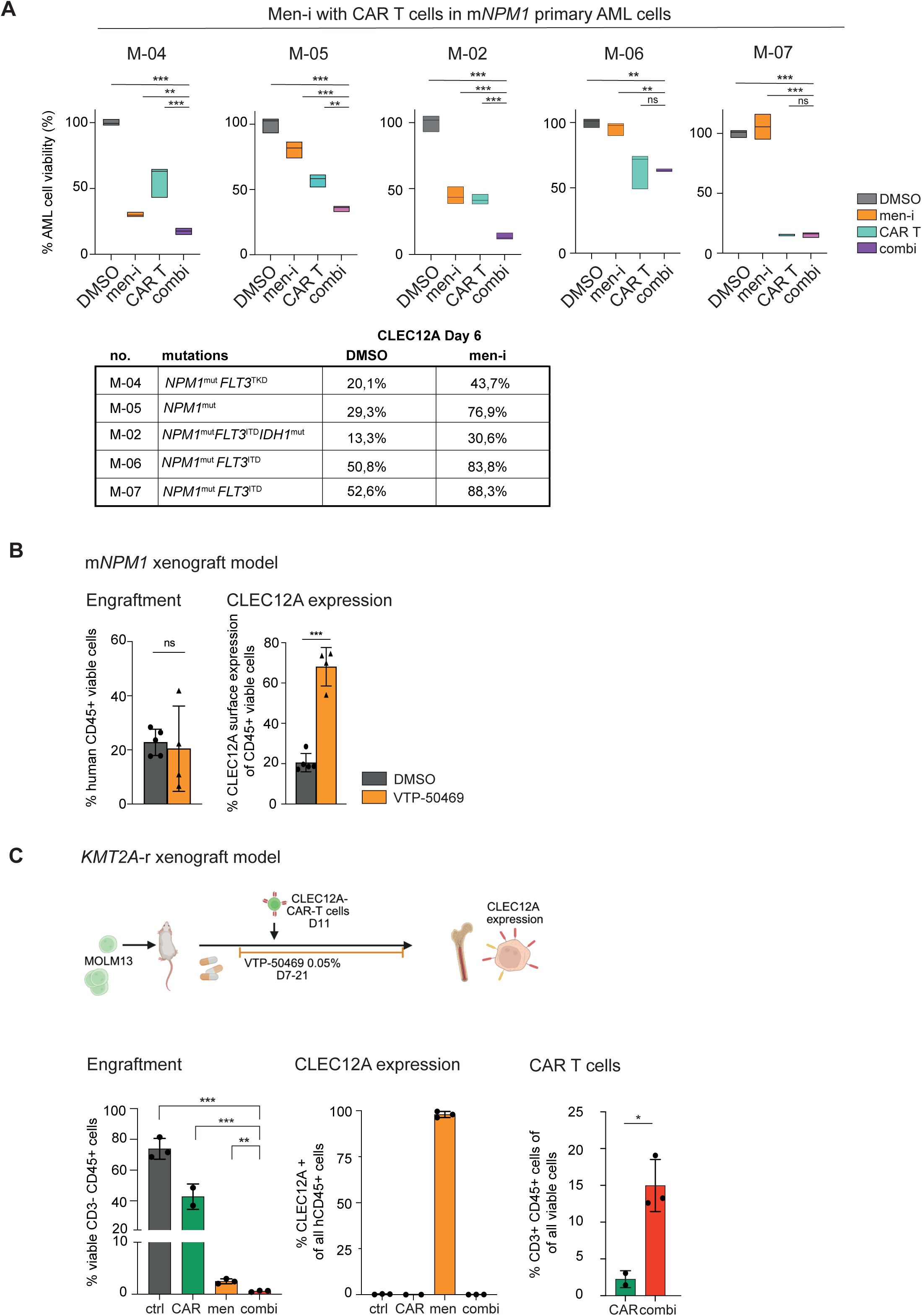
Menin-inhibition with CLEC12A-directed cellular therapy is highly effective against *NPM1*^mut^ primary AML patient cells and significantly reduces leukemic engraftment in *KMT2A*-r AML *in vivo*. **(A) Combining Menin-inhibition with CAR T cells enhances anti-leukemic activity against *NPM1*^mut^ primary AML patient cells.** *NPM1*^mut^ primary AML cells were pretreated for 6 days with menin inhibitor or DMSO. On day 6, cells were counted, stained with CSFE and plated as indicated. Viability was determined by FC after 19h, by assessing the number of CSFE^high^ viable cells relative to the DMSO control group. Displayed is the E:T-ratio of 1:1 for both the combination and single CAR T treatment. Statistical analysis by unpaired t-test, * p<0.05, ** p<0.01, *** p<0.001. **(B) Menin-inhibition increases CLEC12A surface expression in *NPM1*^mut^ AML cells *in vivo.*** OCI-AML3 xenograft model in NSG mice treated with VTP-50469 at 20mg/kg BW bid or vehicle for 14 days. Bone marrow engraftment was determined by FC assessing the CD45+ cells. CLEC12A surface expression was assessed by FC in CD45+CLEC12A+ cells. Statistical analysis by unpaired t-test, *** p <0.001. **(C) Combining menin-inhibition with CLEC12A-directed CAR T cells significantly decreases leukemic bone marrow engraftment in *KMT2A*-r AML *in vivo.*** NSG mice were injected with *KMT2A*-r MOLM13 cells and treated with VTP-chow 0.05% for 14 days, starting on day 7. On day 11, CLEC12A-CAR T cells were injected. Readout was assessed by FC, measuring the human CD3-CD45+ cells compared to all viable cells in the bone marrow on day 21. CLEC12A expression was determined in all human CD45+ cells. CAR T cells were defined as CD3+ CD45+ cells among all viable cells in the bone marrow. Statistical analysis by t-test, * p<0.05, p<0.001. Figure made in BioRender.com.

Our *in vitro* data showed an enhanced anti-leukemic effect when menin-i was combined with CLEC12A-directed CAR T-cell therapy, likely due to menin-induced upregulation of the CLEC12A-CAR target on leukemic blasts. To assess the effects of this combination regimen *in vivo*, we first determined whether CLEC12A could be induced on AML blasts in a *NPM1*^mut^ OCI-AML3 xenograft model, using plasma concentrations typically achieved with standard doses of the menin inhibitor VTP-50469. Therefore, animals were treated for 14 days with VTP-50469 at 40mg/kg/day by oral gavage, then sacrificed, and engrafted human CD45+ AML blasts were assessed for CLEC12A expression compared with vehicle control-treated mice. VTP-50469-treated animals exhibited a 3.3-fold increase in CLEC12A expression compared to the vehicle group (vehicle: 20%; VTP:68% (Fig.6B)).

Having confirmed CLEC12A upregulation on engrafted AML blasts, we next sought to explore the therapeutic potential of combined menin-i and CLEC12A-CAR T-cell therapy *in vivo*. First, we assessed the efficacy of this approach on leukemic burden in a *KMT2A*-r MOLM13-derived AML xenograft model, as these cells exhibit low baseline CLEC12A expression and strong CLEC12A induction upon menin-i (Fig.1C). MOLM13 cells were transplanted into NSG mice via tail-vein injection as described in the methods section, and the animals were randomly divided into 4 groups (vehicle control, VTP, CLEC12A-CAR T, and the combination of VTP and CAR T). As depicted in the schematic in Fig.6C, VTP treatment (0.05% chow) of the single-drug and the combination group animals was initiated on day 7 after transplantation, and the same number of viable CAR T cells was injected into the single CAR and into the combination group animals on day 11. The animals were euthanized on day 21, after 14 days of drug treatment and 10 days after CAR injection. Leukemia burden, as defined by the percentage of bone marrow cells expressing human CD45 but no human CD3 (hCD45+ hCD3-) was significantly reduced within the animal group treated with the VTP and CAR T cell combination compared to all other groups (vehicle: 73.9%±6.75%, CAR: 42.7%%±8.32%; menin-i: 2.5%±0.48%; combination: 0.6%±0.14%, *p*= 0.0029 (combination vs. menin-i) (Fig.6C, Suppl.-Fig.14B). Of note, CLEC12A expression was detected in 97.9% of engrafted hCD45+ AML cells in the single-treatment menin inhibitor group, whereas it was close to 0% in all other groups, including the combination group. These data confirm that CLEC12A was strongly induced by menin-i *in vivo* and are consistent with CAR T cells causing complete eradication of CLEC12A-expressing AML cells in the combination group. It is of interest to note that we detected engrafted CAR T cells, as defined by human CD3-positivity, in 2.25% and 14.98% in the single-CAR vs. combination group, respectively (Fig.6C), consistent with CAR T cell expansion in the latter, possibly due to CLEC12A-target induction on the AML cells.

In a separate experiment, we assessed survival in the disseminated MOLM13 xenograft leukemia model. Drug treatment was initiated on day 7 after transplantation and continued until day 28, and CLEC12A-directed CAR T cells were injected on day 11 (Fig.7A, Suppl.-Fig.14C). Combinatorial VTP and CAR T cell treatment resulted in a significant survival advantage compared with single-drug, single-CAR or vehicle-treated animals (hazard ratios for death 0.5687 (95% confidence interval 0.1422 to 2.274) and 0.1896 (95% confidence interval 0.047-0.758); P=0.0094 and P=0.0006 for combination vs. VTP and combination vs. CAR, respectively; Fig.7B). Notably, the median overall survival was 18 days (vehicle), 20 days (single CAR), 60 days (menin-i) and 99 days (combination). Interestingly, the leukemic blasts of mice treated with the combination of menin-i and CAR T cells showed reduced CLEC12A expression compared to menin inhibitor-treated mice as terminal disease developed (Fig.7C). Cytomorphological analysis revealed signs of differentiation in the AML blasts treated with menin-i (Fig. 7D).

**Figure 7:**
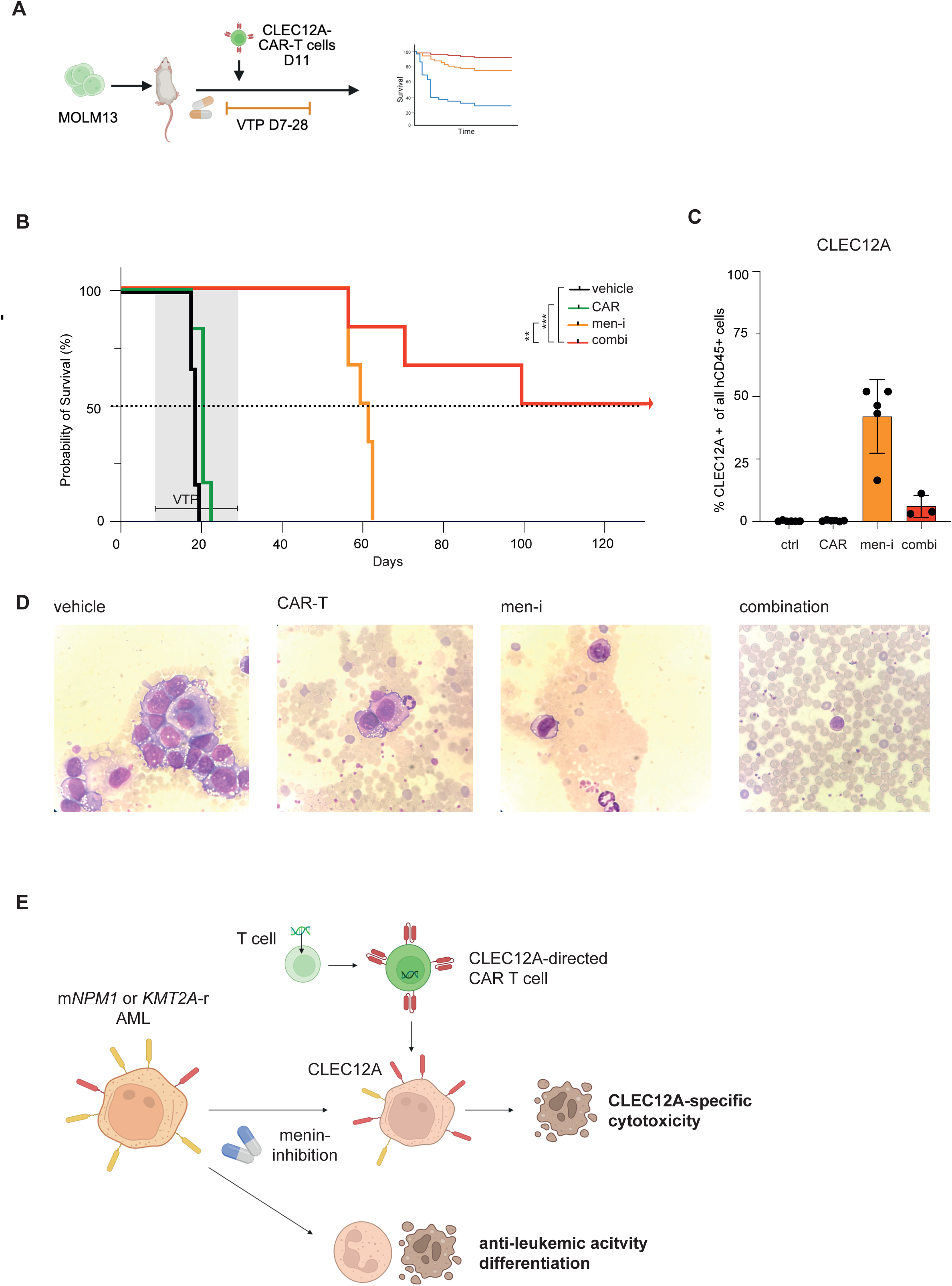
The combination of menin-inhibition with CLEC12A-directed CAR T-cell therapy significantly prolongs the survival against *KMT2A*-r AML *in vivo*. **(A) Experimental Design.** NSG mice were injected with *KMT2A*-r MOLM13 cells and treated with VTP-chow 0.05% for 22 days, starting on day 7. On Day 11, CLEC12A-engineered CAR T cells were injected. Mice were sacrificed when displaying signs of terminal disease. Figure made in BioRender.com. **(B) Combining CLEC12A-directed CAR T-cell therapy with menin-inhibition significantly prolongs the survival of mice in a *KMT2A*-r AML xenograft model.** Treatment as depicted in Fig.7A. Displayed is the overall survival of mice in a Kaplan-Meier diagram. Significance by Log-rank (Mantel-Cox) test. ** p<0.01, *** p<0.001. Data-cutoff was Day 130. **(C) CLEC12A expression at the timepoint of terminal disease.** CLEC12A expression was assessed by FC as CD45+CLEC12A-positive cells of all CD45+ cells. **(D) Representative peripheral blood cytomorphology from mice treated within the vehicle, CAR T, menin inhibitor, or the combination group**. *Picture 1*: AML blasts and murine granulocytes from a mouse of the vehicle group. *Picture 2*: AML blasts from a mouse treated with VTP-50469 (at terminal disease), consistent with signs of monocytic differentiation. *Picture 3*: CAR T cell in the peripheral blood of a mouse treated with the combination of menin-i and CLEC12A-CAR T cells. **(E) Proposal of the therapy model.** Menin-i has strong anti-leukemic activity and induces differentiation in *KMT2A*-r and *NPM1*^mut^ AMLs. It also upregulates the immune target CLEC12A. This can be targeted by CLEC12A-directed CAR T cells, resulting in target-specific AML killing. Together, the combination of menin-i and CLEC12A-directed CAR T cells significantly improves anti-leukemic activity compared with single treatments *in vitro* and *in vivo.* Figure made in BioRender.com.

In summary, these data confirm that the combination of menin-i with CLEC12A-directed CAR T cells substantially improves survival of mice engrafted with *KMT2A*-r AML compared with single-drug, single-CAR, or vehicle-treated animals, and support this therapeutic concept as a synergistic approach to combat these leukemias (Fig.7E).

## Discussion

Menin inhibitors are the first specific targeted agents approved against *KMT2A*-r and the most common *NPM1*^mut^ AML subtype. These novel inhibitors have demonstrated remarkable activity in early clinical trials against relapsed or refractory disease and represent one of the most consequential advances in leukemia treatment (reviewed in (36),(12)). Despite their astonishing ability to induce remissions in heavily pretreated leukemia patients (13–16), the majority of patients treated with single-agent menin-i eventually develop disease progression (37). Combination therapy has the potential to overcome these challenges, and synergistic menin inhibitor combinations from preclinical studies with FLT3-, BCL2-, PRMT5-, DOT1L-inhibitors, or IMiDs are awaiting clinical validation (4,5,30,31,38,39).

Cancer immunotherapy, and specifically CAR-based cellular therapy, has reshaped the treatment of lymphoid neoplasms in recent years, leading to deep and durable responses (17–21). CAR-based cellular therapy against AML, however, has shown low efficacy in clinical studies to date (reviewed in (22)). The majority of clinical trials assessing CARs in AML have targeted CD33, CD123, and CLEC12A. Small clinical trials and case series assessing CD33- and CD123-directed CARs in AML reported only a few responses and commonly resulted in marrow aplasia. Although CLEC12A-directed CAR treatment was associated with higher CR rates, those were commonly followed by early relapses if no subsequent allo-SCT was performed (26–29). Potential explanations include heterogeneous CLEC12A expression and variable antigen density in AML-initiating cell populations (22,24,40).

In this study, we show that menin inhibitors can be utilized to prime AML cells for a CLEC12A-directed CAR therapy by uniformly inducing strong transcriptional CLEC12A upregulation that translates into broad CLEC12A cell-surface expression in *KMT2A*-r and *NPM1*^mut^ AML (Fig.1A-D, Suppl-Fig.1A-C). Given that CLEC12A is not bound by the menin-KMT2A complex before or after menin-i, CLEC12A is likely not induced via a direct chromatin-based transcriptional activation mechanism (Suppl.-Fig.1D,E). As menin-i induces strong monocytic/neutrophilic AML cell differentiation and since CLEC12A is normally expressed on myelo-monocytic cells, including neutrophils and monocytes (25), CLEC12A-induction may be caused by menin inhibitor-associated differentiation processes in the AML blasts. Notably, ATRA-induced differentiation of AML cell lines into neutrophils did not induce CLEC12A (Suppl.-Fig.2A-C), and further studies will be necessary to determine whether CLEC12A induction is menin inhibitor-specific or also induced by other drugs. The concept of using epigenetic targeting to sensitize AML to immunotherapy, however, has been proposed before. DNA hypomethylating agents (HMAs), such as decitabine or azacitidine, have been shown to reverse MHC-II expression loss after allo-SCT and are currently used in high-risk post-allo-SCT settings to prevent or treat AML relapse, with moderate success (41,42). While a recent preclinical study demonstrated that menin-i upregulates MHC-II molecules and thereby enhances the graft-versus-leukemia effect in a human xenograft transplantation model (43), our study is, to our knowledge, the first example of a target-selective epigenetic compound that uniformly upregulates a specific immunostimulatory antigen for CAR-directed cellular therapy. Furthermore, in our non-genetic (*MEN1*-wildtype) menin inhibitor-resistance model of *NPM1*^mut^ AML, CLEC12A induction was maintained (Suppl.Fig.1C), leaving those cells potentially prone to CLEC12A-directed targeting even in the presence of menin inhibitor resistance. While non-genetic resistance mechanisms may very well be the dominant mechanism of menin inhibitor resistance, according to a recent study assessing AML patients who developed menin inhibitor resistance (44), it remains to be seen whether CLEC12A may also be maintained in AML patients, progressing on menin-i.

In addition, menin-i not only induces CLEC12A, but also dramatically enhances the anti-leukemic efficacy of CLEC12A-engineered CAR therapy against AML dramatically when used in combination (Fig.5, Fig.6A, Suppl-Fig.11-13) and independently of CLEC12A baseline expression. While CLEC12A is already one of the more promising antigen targets for CAR therapy in AML, rapid relapses emphasize the need for further improvement.

Menin inhibitors are - unlike chemotherapy or other immunosuppressing agents - particularly attractive combination partners for cellular immunotherapy, as they do not negatively affect immune effector cell functionality but have strong anti-leukemic effects. While few studies have so far investigated menin inhibitor effects on T and NK cells, we characterized the effects of VTP-50469 in detail on those cells and found no significant inhibitory effects on gene expression, immunophenotype, IFN-γ release, and killing efficacy (Fig.2, Suppl.-Fig.3-5). Only extended 21-day exposure to very high concentrations of the drug had minor anti-proliferative effects on cultured T cells, while NK cells continued to proliferate even at a slightly increased pace (Fig.2C). Furthermore, single-cell transcriptomic profiling of T and NK cells showed that menin-i does not measurably impact the assembly of transcriptionally defined subpopulations and revealed no differentially expressed genes within subclusters indicative of senescence or functional impairment (Fig.3, Suppl.-Fig.6-9). While these data support the concept of combining menin-i with CAR-based immunotherapy, they may also be of interest in the context of post-allo-SCT clinical situations and suggest that potential menin inhibitor-maintenance therapy is unlikely to negatively influence the functionality of engrafted immune effector cells. These data are therefore consistent with a recent publication demonstrating that menin-i may even enhance the graft-versus-leukemia effect following allo-SCT under specific conditions (43).

It is of interest to note that menin inhibitors appear to have a slight positive effect on NK cell proliferation and IFN-γ response (Fig. 2C,E), thereby advocating menin inhibitor combinations with CAR NK cells. Since the CLEC12A-directed CAR NK cells only show slightly enhanced killing vs. NT NK cells in our *in vitro* assays - most likely due to their very strong alloreactive effects - and some potential fratricide activity due to low CLEC12A NK cell self-expression, we focused in this study on the *in vivo* assessment with CAR T cells. However, given the potential alloreactivity against AML even in the case of target-antigen loss (35,45,46), and the lack of a CAR NK cell memory effect that may prevent long-term hematotoxicity, this approach warrants further investigation in future studies.

A particularly exciting aspect of this study is the remarkable activity of combined menin-i with CLEC12A-directed CAR T-cell therapy in a very aggressive human AML xenograft model with almost no CLEC12A baseline expression. While all animals in the vehicle control, the CAR T cell, and in the VTP-50469 monotherapy group died of AML within 18, 20, and 60 days, respectively, survival of the animals treated with combined menin-i and CLEC12A CAR T cells was dramatically longer with half of the animals still being alive and healthy after 130 days with no engrafted leukemia cells in the peripheral blood, likely consistent with eradicated disease (Fig.7). Only 14 days of continuous VTP-50469 treatment at a low-effective dose (0.05%) efficiently induced CLEC12A expression, enabling efficient CAR-directed targeting within the combinatorial group, with no clinically apparent adverse effects. The astonishing efficacy of combined menin-i with CLEC12A-directed CAR effector-cell therapy calls for clinical validation in human trials and is readily amenable to translation into early-phase human trials.

## Material and Methods

### Cell culture of cell lines

All human cell lines used in this study were authenticated by Multiplex Authentication by Multiplexion (Heidelberg, Germany) between July 2022 and July 2024 as previously described (5), or obtained in early passage from the Leibniz Institute DSMZ. Cells were maintained under standard conditions (4,5) as described in the supplemental methods.

### Cell culture of immune cells

Primary T and NK cells were isolated from peripheral blood mononuclear cells (PBMCs) from healthy donors and cultivated in respective T or NK cell media. For long term cultivation, T cells were either re-stimulated by the addition of lethally irradiated OCI-AML3 (150.000 cells/ml) or agonistic anti-CD3/CD28 DynaBeads and 50-200IU/mL recombinant IL-2. Primary NK cells were stimulated with 120 IU/mL IL-15. Purity and viability of isolated T and NK cells was confirmed by FC (FACS Celesta or FACS Canto II, BD). Detailed information is described in the supplemental methods.

### In vitro studies

*In vitro* drug treatment, cell viability assays, FC-based surface marker assessment, ELISpot assays, ChIP sequencing and morphological analysis were performed according to standard procedures with detailed description in the supplemental methods section.

### Human Primary AML Blast Assay

Primary AML samples were obtained under Institutional Review-Board approved protocols from patients diagnosed at the University Medical Center Mainz or Frankfurt, following written consent. Primary AML cells were cultured in serum-free media supplemented with different cytokines as previously described (5,47) and provided in the supplemental methods. Cells were exposed for six days to 100nM VTP-50469 or DMSO before assessing CLEC12A surface expression and combined anti-leukemic efficacy of menin-i and CAR T-cell therapy.

### Single Cell Sequencing

CD3+ T and CD56+ NK cells from four different healthy donors were cultured in lineage-specific media supplemented with 50IU/mL IL-2 and agonistic anti-CD3/CD28 DynaBeads (T cells) or 120IU/ml IL-15 (NK cells) for 7 days. Cells were then treated with 100nM VTP-50469 or DMSO-control for an additional 4 days, washed on day 4 to increase viability, and processed for single-cell sequencing using the BD Rhapsody (platform BD Biosciences) with 2,000 cells per sample at ∼30,000 reads per cell. Four biological replicates were analyzed per group. Detailed bioinformatical analyses are described in the supplemental methods.

### CLEC12A-CAR design and production of lentiviral vectors

The second-generation, 4-1BB-based CAR sequence *C12A.BBz* was designed *in silico*, followed by integration into the third-generation lentiviral transfer plasmid pSL-GG, as described in the supplemental methods. VSV-G- and RD114TR-pseudotyped, self-inactivating lentiviral vectors (LV) were produced using a third-generation plasmid system by polyethyleneimine (PEI)-based (Polysciences) transfection of Lenti-X™ 293T cells (Takara Bio). Detailed description of the virus production is provided in the supplemental methods.

### Transduction of primary immune cells

Activated T cells were transduced with VSV-G-pseudotyped LV vectors (MOI = 1) diluted in RPMI 1640 by spin-infection at 800xg for 90 minutes at 32°C, as recently described (48). Transduction of primary NK cells (MOI=1) was performed on day two post isolation as recently described (49,50). CAR expression and phenotypes of engineered CAR T and CAR NK cells were monitored by FC (FACS Celesta & FACS Canto II, BD). A detailed description is provided in the supplemental methods.

### *In vitro* Cytotoxicity Assays

Cytotoxic functionality of immune cells was assessed in FC-based and luciferase-based assays against the AML cells lines OCI-AML2, OCI-AML3, MOLM13 or primary AML cells. Tumor cells expressed luciferase (*FLUC*-OCI-AML3) or were labeled with CFSE (Thermo Fisher Scientific) according to the manufacturer’s instructions before co-culture with CAR or non-transduced immune cells at indicated E:T-ratios. AML cells without effector cells (AML only) served as control for spontaneous lysis. Viability was assessed after 4 to 24h, estimated by omitted luminescence (plate reader, FluOstar OMEGA 0415, 2007) or viable CSFE^high^DAPI^low^ cells (FC, FACS CANTO II, BD) normalized to the control group (AML only). Independent experiments were conducted in technical triplicates with immune cells from different biological donors. A detailed method and data analysis description is provided in the supplemental methods section.

### *In vitro* Combination Assays

Target cells (MOLM13, OCI-AML3 or primary AML cells) were pretreated with 100nM VTP-50469 (MedChem Express) or DMSO for four (cell lines) or six (primary cells) days and were labeled as previously described. An equal number of viable target cells per well was plated for either condition (menin-i or DMSO) on day four and immune cells added in different E:T-ratios (3:1 to 1:27) in lineage-specific media. Viability was determined after 6 or 19h normalizing the respective luminescence (plate reader, FluOstar OMEGA 0415, 2007) or viable CSFE^high^DAPI^low^ cells (FC) to the luminescence/number of DMSO-pretreated control cells (DMSO only).

### In vivo Experiments

1×10^6^ MOLM13 cells were injected into 8-15 week-old NOD.Cg-*Prkdc^scid^ Il2rg^tm1Wjl^*/SzJ (NSG) mice. Animals were randomized into four treatment groups: vehicle, VTP-chow 0.05%, CLEC12A-directed CAR T cells (1×10^7^/mouse) and their combination. Treatment with VTP-chow started on day 7 and was continued for 14 days (until day 21) for the leukemic burden experiment, or 22 days (until day 29) for the survival experiment. CAR T cells were tail vein injected on day 11. For the assessment of leukemia burden, the mice were euthanized, the bone marrow harvested and leukemic engraftment determined by measuring the viable human CD45+CD3-cells using FC on day 21. For survival analysis, moribund mice were euthanized at signs of terminal disease. *In vivo* experiments were approved by the National Investigation Offices of Mainz, Rheinland-Pfalz and Freiburg, Baden-Württemberg.

### Data Analysis and Statistical Methods

Statistical test computations were performed using Graph Pad Prism (v10). Statistical significance for the *in vitro* assays was calculated using the Student’s *t*-test. Survival analysis was estimated using the Kaplan-Meier method, and *p-*values were calculated using the log-rank (Mantel-Cox) test. *P-*values were displayed as follows: *, *P* < 0.05; **, *P* < 0.01; ***, *P* < 0.001. All *in vitro* data represent the mean of at least three independent experiments. Immunophenotyping assays were performed in technical duplicates, all other *in vitro* assays in technical triplicates. Experiments involving immune cells used different biological donors for the independent experiments.

## Data Availability Statement

The data generated in this study are available upon request from the corresponding author. Other data generated in this study are available within the article and its supplementary data files. Schematic figures were created with BioRender.com (Fig.2A, Fig.3A, Fig.4A+B, Fig.5A, Fig.6C, Fig.7A+E, and Suppl.Fig.5E).

## Acknowledgments

The authors thank Annkatrin Gebel for expert technical assistance; the Forschungszentrum for Immuntherapie (FZI) for assisting with single cell sequencing and bioinformatic analyses; the Translational Animal Research Center (TARC) of the University Medical Center, Mainz, for providing the animals, the team of the Biobank at the UCT Frankfurt (under excellent guidance of T. Oellerich and C. Brandts) for support with primary patient samples, and Basepair for bioinformatics support.

This work was funded by grants from the Deutsche Forschungsgemeinschaft (DFG) to MWMK: KU-2688/2-1 and KU-2688/2-2 as well as part of the Collaborative Research Center (CRC) 1292 (SFB1292/2/TP12 and SFB1292/3/TP12; to MWMK and EU). This work was in part supported by the Stiftung Deutsche Krebshilfe (German Cancer Aid): consortium “CAR FACTORY” #70115200 as part of the preclinical cancer drug development network (preCDD; to EU), by the German Cancer Consortium (DKTK) and German Cancer Research Center (DKFZ; to EU, PW, LRK and RZ), and the Else Kröner-Fresenius-Stiftung (EKFS; to EU).

RZ was supported by SFB-1479 (Project ID: 441891347, P01; to R.Z.), by the DFG (RU5659 TARGET-MPN: ZE 872/6-1, TP 7; to RZ), the European Union (EU Proposal n°ERC-2022-ADG Project: 101094168; to RZ), by AlloCure (ERC Advanced grant; to R.Z.), and the Germany’s Excellence Strategy (CIBSS – EXC-2189 – Project ID 390939984; to R.Z.). The data published in this work is part of the scientific dissertation of Philipp Wendel (PW) and the medical dissertation from Marie Kuhmann (MK) and Lilian Viehböck (LV).

## Authorship Contributions

MWMK designed the research, analyzed and interpreted the data, wrote and revised the manuscript and supervised the study. EU designed the research, analyzed and interpreted the data and revised the manuscript. RZ analyzed and interpreted the data and revised the manuscript. JR and PW performed the experiments, analyzed and interpreted data, wrote the original draft of the manuscript and edited the manuscript. MD, MS, FG, VF, LRK, MK, LV, SW, ND, JH, CL, MK and CW performed experiments, analyzed and interpreted data. NA, FM and MD analyzed and interpreted bioinformatic data. KD and HD characterized and provided primary AML patient samples and revised the manuscript. CW, HE, MT, and DS revised the manuscript.

## References

1. Kühn MWM, Pemmaraju N, Heidel FH. The evolving landscape of epigenetic target molecules and therapies in myeloid cancers: focus on acute myeloid leukemia and myeloproliferative neoplasms. Leukemia. 2025;39:1824–37.

2. Borkin D, He S, Miao H, Kempinska K, Pollock J, Chase J, et al. Pharmacologic Inhibition of the Menin-MLL Interaction Blocks Progression of MLL Leukemia In Vivo. Cancer Cell. 2015;27:589–602.

3. Yokoyama A, Somervaille TCP, Smith KS, Rozenblatt-Rosen O, Meyerson M, Cleary ML. The Menin Tumor Suppressor Protein Is an Essential Oncogenic Cofactor for MLL-Associated Leukemogenesis. Cell. 2005;123:207–18.

4. Kühn MWM, Song E, Feng Z, Sinha A, Chen C-W, Deshpande AJ, et al. Targeting Chromatin Regulators Inhibits Leukemogenic Gene Expression in NPM1 Mutant Leukemia. Cancer Discov. 2016;6:1166–81.

5. Dzama MM, Steiner M, Rausch J, Sasca D, Schönfeld J, Kunz K, et al. Synergistic targeting of FLT3 mutations in AML via combined menin-MLL and FLT3 inhibition. Blood. 2020;136:2442–56.

6. Wang XQD, Fan D, Han Q, Liu Y, Miao H, Wang X, et al. Mutant NPM1 Hijacks Transcriptional Hubs to Maintain Pathogenic Gene Programs in Acute Myeloid Leukemia. Cancer Discov. 2023;13:724–45.

7. Uckelmann HJ, Haarer EL, Takeda R, Wong EM, Hatton C, Marinaccio C, et al. Mutant NPM1 Directly Regulates Oncogenic Transcription in Acute Myeloid Leukemia. Cancer Discov. 2023;13:746–65.

8. Klossowski S, Miao H, Kempinska K, Wu T, Purohit T, Kim E, et al. Menin inhibitor MI-3454 induces remission in MLL1-rearranged and NPM1-mutated models of leukemia. J Clin Invest. 130:981–97.

9. Kwon MC, Thuring JW, Querolle O, Dai X, Verhulst T, Pande V, et al. Preclinical efficacy of the potent, selective menin-KMT2A inhibitor JNJ-75276617 (bleximenib) in KMT2A- and NPM1-altered leukemias. Blood. 2024;144:1206–20.

10. Krivtsov AV, Evans K, Gadrey JY, Eschle BK, Hatton C, Uckelmann HJ, et al. A Menin-MLL Inhibitor Induces Specific Chromatin Changes and Eradicates Disease in Models of MLL-Rearranged Leukemia. Cancer Cell. 2019;36:660–673.e11.

11. Uckelmann HJ, Kim SM, Wong EM, Hatton C, Giovinazzo H, Gadrey JY, et al. Therapeutic targeting of preleukemia cells in a mouse model of NPM1 mutant acute myeloid leukemia. Science; 2020;367:586–90.

12. Kühn MWM, Ganser A. The Menin story in acute myeloid leukaemia-The road to success. Br J Haematol. 2024;205:812–4.

13. Wang ES, Issa GC, Erba HP, Altman JK, Montesinos P, DeBotton S, et al. Ziftomenib in relapsed or refractory acute myeloid leukaemia (KOMET-001): a multicentre, open-label, multi-cohort, phase 1 trial. Lancet Oncol. 2024;25:1310–24.

14. Issa GC, Aldoss I, DiPersio J, Cuglievan B, Stone R, Arellano M, et al. The menin inhibitor revumenib in KMT2A-rearranged or NPM1-mutant leukaemia. Nature. 2023;615:920–4.

15. Issa GC, Aldoss I, Thirman MJ, DiPersio J, Arellano M, Blachly JS, et al. Menin Inhibition With Revumenib for KMT2A-Rearranged Relapsed or Refractory Acute Leukemia (AUGMENT-101). J Clin Oncol. 2025;43:75–84.

16. Arellano ML, Thirman MJ, DiPersio JF, Heiblig M, Stein EM, Schuh AC, et al. Menin inhibition with revumenib for NPM1-mutated relapsed or refractory acute myeloid leukemia: the AUGMENT-101 study. Blood. 2025;146:1065–77.

17. Rodriguez-Otero P, Ailawadhi S, Arnulf B, Patel K, Cavo M, Nooka AK, et al. Ide-cel or Standard Regimens in Relapsed and Refractory Multiple Myeloma. New England Journal of Medicine. 2023;388:1002–14.

18. Martin T, Usmani SZ, Berdeja JG, Agha M, Cohen AD, Hari P, et al. Ciltacabtagene Autoleucel, an Anti-B-cell Maturation Antigen Chimeric Antigen Receptor T-Cell Therapy, for Relapsed/Refractory Multiple Myeloma: CARTITUDE-1 2-Year Follow-Up. J Clin Oncol. 2023;41:1265–74.

19. Schuster SJ, Bishop MR, Tam CS, Waller EK, Borchmann P, McGuirk JP, et al. Tisagenlecleucel in Adult Relapsed or Refractory Diffuse Large B-Cell Lymphoma. New England Journal of Medicine. 2019;380:45–56.

20. Neelapu SS, Locke FL, Bartlett NL, Lekakis LJ, Miklos DB, Jacobson CA, et al. Axicabtagene Ciloleucel CAR T-Cell Therapy in Refractory Large B-Cell Lymphoma. N England Journal of Medicine. 2017;377:2531–44.

21. Cappell KM, Kochenderfer JN. Long-term outcomes following CAR T cell therapy: what we know so far. Nat Rev Clin Oncol. 2023;20:359–71.

22. Haubner S, Subklewe M, Sadelain M. Honing CAR T cells to tackle acute myeloid leukemia. Blood. 2025;145:1113–25.

23. van Rhenen A, van Dongen GAMS, Kelder A, Rombouts EJ, Feller N, Moshaver B, et al. The novel AML stem cell–associated antigen CLL-1 aids in discrimination between normal and leukemic stem cells. Blood. 2007;110:2659–66.

24. Haubner S, Mansilla-Soto J, Nataraj S, Kogel F, Chang Q, de Stanchina E, et al. Cooperative CAR targeting to selectively eliminate AML and minimize escape. Cancer Cell. 2023;41:1871–1891.e6.

25. Marshall ASJ, Willment JA, Lin H-H, Williams DL, Gordon S, Brown GD. Identification and characterization of a novel human myeloid inhibitory C-type lectin-like receptor (MICL) that is predominantly expressed on granulocytes and monocytes. J Biol Chem. 2004;279:14792–802.

26. Ma Y-J, Dai H-P, Cui Q-Y, Cui W, Zhu W-J, Qu C-J, et al. Successful application of PD-1 knockdown CLL-1 CAR-T therapy in two AML patients with post-transplant relapse and failure of anti-CD38 CAR-T cell treatment. Am J Cancer Res. 2022;12:615–21.

27. Zhao Y, Zhang X, Zhang M, Guo R, Zhang Y, Pu Y, et al. Modified EASIX scores predict severe CRS/ICANS in patients with acute myeloid leukemia following CLL1 CAR-T cell therapy. Ann Hematol. 2024;103:969–80.

28. Zhang H, Bu C, Peng Z, Li G, Zhou Z, Ding W, et al. Characteristics of anti-CLL1 based CAR-T therapy for children with relapsed or refractory acute myeloid leukemia: the multi-center efficacy and safety interim analysis. Leukemia. 2022;36:2596–604.

29. Pei K, Xu H, Wang P, Gan W, Hu Z, Su X, et al. Anti-CLL1-based CAR T-cells with 4-1-BB or CD28/CD27 stimulatory domains in treating childhood refractory/relapsed acute myeloid leukemia. Cancer Med. 2023;12:9655–61.

30. Aubrey BJ, Cutler JA, Bourgeois W, Donovan KA, Gu S, Hatton C, et al. IKAROS and MENIN coordinate therapeutically actionable leukemogenic gene expression in MLL-r acute myeloid leukemia. Nat Cancer. 2022;3:595–613.

31. Rausch J, Dzama MM, Dolgikh N, Stiller HL, Bohl SR, Lahrmann C, et al. Menin inhibitor ziftomenib (KO-539) synergizes with drugs targeting chromatin regulation or apoptosis and sensitizes acute myeloid leukemia with MLL rearrangement or NPM1 mutation to venetoclax. Haematologica. 2023;108(10):2837–2843

32. Uhlén M, Fagerberg L, Hallström BM, Lindskog C, Oksvold P, Mardinoglu A, et al. Proteomics. Tissue-based map of the human proteome. Science. 2015;347:1260419.

33. Gíslason MH, Demircan GS, Prachar M, Furtwängler B, Schwaller J, Schoof EM, et al. BloodSpot 3.0: a database of gene and protein expression data in normal and malignant haematopoiesis. Nucleic Acids Research. 2024;52:D1138–42.

34. Rebuffet L, Melsen JE, Escalière B, Basurto-Lozada D, Bhandoola A, Björkström NK, et al. High-dimensional single-cell analysis of human natural killer cell heterogeneity. Nat Immunol. 2024;25:1474–88.

35. Albinger N, Pfeifer R, Nitsche M, Mertlitz S, Campe J, Stein K, et al. Primary CD33-targeting CAR-NK cells for the treatment of acute myeloid leukemia. Blood Cancer J. 2022;12:61.

36. Seeling C, Ganser A, Döhner H, Kühn MWM. Tailoring intensive and less-intensive treatment in acute myeloid leukemia. Semin Hematol. 2025;62:196–208.

37. Chin K-K, Ball BJ, Abaza Y, Altman JK, Thakur RK, Ali M, et al. Outcomes of relapsed or refractory acute myeloid leukemia after menin inhibition failure. Blood Adv. 2025; bloodadvances.2025018178.

38. Fiskus W, Boettcher S, Daver N, Mill CP, Sasaki K, Birdwell CE, et al. Effective Menin inhibitor-based combinations against AML with MLL rearrangement or NPM1 mutation (NPM1c). Blood Cancer J. 2022;12:1–11.

39. Carter BZ, Tao W, Mak PY, Ostermann LB, Mak D, McGeehan G, et al. Menin inhibition decreases Bcl-2 and synergizes with venetoclax in NPM1/FLT3-mutated AML. Blood. 2021;138:1637–41.

40. Willier S, Rothämel P, Hastreiter M, Wilhelm J, Stenger D, Blaeschke F, et al. CLEC12A and CD33 coexpression as a preferential target for pediatric AML combinatorial immunotherapy. Blood. 2021;137:1037–49.

41. Christopher MJ, Petti AA, Rettig MP, Miller CA, Chendamarai E, Duncavage EJ, et al. Immune Escape of Relapsed AML Cells after Allogeneic Transplantation. New England Journal of Medicine. 2018;379:2330–41.

42. Craddock C, Labopin M, Robin M, Finke J, Chevallier P, Yakoub-Agha I, et al. Clinical activity of azacitidine in patients who relapse after allogeneic stem cell transplantation for acute myeloid leukemia. Haematologica. 2016;101:879–83.

43. Fetsch V, Schwöbel LF, Ozyerli-Goknar E, Stell A-V, Punta M, Plenge T, et al. Menin inhibition enhances graft-versus-leukemia effects by T-cell activation and endogenous retrovirus induction in AML. Blood. 2026 Jan 29;147(5):584–601.

44. Gordon SJV, Perner F, MacPherson L, Fennell KA, Wenge DV, Bourgeois W, et al. Catalytic Inhibition of KAT6/KAT7 Enhances the Efficacy and Overcomes Primary and Acquired Resistance to Menin Inhibitors in MLL Leukemia. Cancer Discov. 2025;15:2117–38.

45. Laskowski TJ, Biederstädt A, Rezvani K. Natural killer cells in antitumour adoptive cell immunotherapy. Nat Rev Cancer. 2022;22:557–75.

46. Liu E, Marin D, Banerjee P, Macapinlac HA, Thompson P, Basar R, et al. Use of CAR-Transduced Natural Killer Cells in CD19-Positive Lymphoid Tumors. N England Journal of Medicine. 2020;382:545–53.

47. Kühn MWM, Hadler MJ, Daigle SR, Koche RP, Krivtsov AV, Olhava EJ, et al. MLL partial tandem duplication leukemia cells are sensitive to small molecule DOT1L inhibition. Haematologica. 2015;100:e190–3.

48. Prommersberger S, Hudecek M, Nerreter T. Antibody-Based CAR T Cells Produced by Lentiviral Transduction. Curr Protoc Immunol. 2020;128:e93.

49. Bexte T, Albinger N, Al Ajami A, Wendel P, Buchinger L, Gessner A, et al. CRISPR/Cas9 editing of NKG2A improves the efficacy of primary CD33-directed chimeric antigen receptor natural killer cells. Nat Commun. 2024;15:8439.

50. Reindl LM, Jalili L, Bexte T, Harenkamp S, Thul S, Hehlgans S, et al. Precision targeting of rhabdomyosarcoma by combining primary CAR NK cells and radiotherapy. J Immunother Cancer. 2025;13:e011330.

