## Supplemental Methods + Figures for "Menin-Inhibition Sensitizes Acute Myeloid Leukemia to CLEC12A-Directed CAR Cell Therapy"

[1]Department of Hematology and Medical Oncology, University Medical Center, Johannes Gutenberg University, Mainz, Germany
[2]German Cancer Consortium (DKTK), Partner site Frankfurt/Mainz and German Cancer Research Center (DKFZ), Heidelberg, Germany
[3]Goethe University, Department of Pediatrics, Experimental Immunology and Cell Therapy, Frankfurt am Main, Germany
[4]Institute for Organic Chemistry and Biochemistry, Technical University of Darmstadt, Darmstadt, Germany
[5]Frankfurt Cancer Institute (FCI), Goethe University, Frankfurt am Main, Germany
[6]Department of Medicine I, Medical Center - University of Freiburg, Faculty of Medicine, University of Freiburg, Freiburg, Germany
[7]Faculty of Biology, Albert-Ludwigs-University, Freiburg, Germany
[8]German Cancer Consortium (DKTK), Partner site Freiburg and German Cancer Research Center (DKFZ), Heidelberg, Germany
[9]Institute of Medical Biostatistics, Epidemiology and Informatics (IMBEI), University Medical Center of the Johannes Gutenberg University Mainz, Mainz, Germany
[10]Research Center for Immunotherapy (FZI), University Medical Center Mainz, Mainz, Germany
[11]Institute of Immunology, University Medical Center, Johannes Gutenberg University, Mainz, Germany
[12]Department of Internal Medicine III, Ulm University Hospital, Ulm, Germany
*co-first authors; #co-senior authors

#### Supplemental Methods

##### Cell culture of cell lines

The human AML cell lines MV411, MOLM13, OCI-AML3, NB4 and HL60 were cultured in RPMI 1640 media (Gibco) supplemented with 10% fetal bovine serum (FBS, BioTrend), 1% Penicillin/Streptomycin (P/S, Sigma), and 1% L-Glutamine (L-Glu, Sigma). OCI-AML2 cells were cultured in MEM α supplemented with nucleosides, GlutaMAX (all Gibco), 20% FBS (BioTrend or Thermo fisher) and 1% P/S (Gibco).

##### Adaptive resistance model in OCI-AML3

OCI-AML3 cells were cultivated for one month in methylcellulose in the presence of the menin inhibitor MI-503 at the $IC_{20}$ value of these cells. Three individual blast colonies, each derived from a single cell, were picked to generate three independent menin inhibitor resistant cell lines. Over a period of five months, the cells were expanded in increasing concentrations of the menin inhibitor MI-503. The cells did not acquire any known resistance mutations in the menin binding pocket (bulk sequencing, (not shown)) and show a cross-resistance against bleximenib, ziftomenib and VTP-50469 (close analogue to revumenib).

##### Isolation and cell culture of immune cells

Ethical approval was granted (approval 329/10 & 274/18), and the blood donated after written consent and in accordance with the Declaration of Helsinki. Peripheral blood mononuclear cells (PBMCs) were isolated from buffy coats of healthy donors provided by the German Red Cross Blood Donation Service Baden-Württemberg-Hessen (Frankfurt am Main, Germany) and the Transfusion Center of the University Medical Center of the Johannes Gutenberg-University (Mainz, Germany), using density gradient centrifugation. Primary T cells were enriched using the EasySep Human CD3 Positive Selection Kit II or EasySep Human T Cell Enrichment Kit EasySep™ (both STEMCELL Technologies) according to the manufacturer's instructions. Primary T cells were cultivated in T cell media (TCMp) composed of RPMI 1640 GlutaMAX supplemented with 10% (v/v) heat-inactivated human plasma type AB (DRK-Blutspendedienst) or heat-inactivated human plasma type B (Transfusion Center, University Medical Center of the Johannes Gutenberg-University Mainz, Germany), 25 mM HEPES (Sigma Aldrich) and 1% (v/v) P/S (Gibco), and 1% L-Glu (Sigma) supplemented with 50 or 200 IU/ml IL-2 (PeproTech) depending on the assay. Cultivation of the T cells was performed at a cell density of $0.5x10^6$ cells/mL with media change every three to four days. Non-transduced (NT) T cells were either re-stimulated by the addition of lethally irradiated OCI-AML3 (150.000/ml) or agonistic anti-CD3/CD28 DynaBeads (ratio 0.1-0.2:1) (Gibco) every seven days. Primary NK cells were enriched using the EasySep™ Human NK Cell Enrichment Kit (STEMCELL Technologies) according to the manufacturer's instructions and cultivated in NK MACS supplemented with NK MACS supplements (both Miltenyi Biotech), 5% (v/v) heat-inactivated human plasma type AB, 1% (v/v) P/S, NK MACS supplements and 120 IU/mL IL-15 (Miltenyi Biotech or PeproTech), as recently described (1).

##### Small molecule inhibitors

VTP-50469, All-trans Retinoid Acid (ATRA) and VTP Fumerate were purchased by MedChem Express. For *in vitro* studies, drugs were dissolved in DMSO and stored at -20°C. For *in vivo* studies, VTP fumerate was pressed in chow by Ssniff Spezialdiäten at a concentration of 0.05%.

##### *In vitro* cell viability assays

20,000 Trypan-negative AML cells or 30,000 NK or T cells per well were plated in 150µL media in 96-well plates. Cells were treated with VTP-50469 in a serial dilution of 9 concentrations (max. concentration 100nM) and DMSO control in technical triplicates. Cell viability was assessed on day 7,14, and 21 assessing DAPI negative cells (Roth, 1µg/mL) by flow cytometry. Cells were split and reseeded every 3-4 days at a density of 20,000 cells or 30,000 cells per well, respectively. For T cell proliferation assays with irradiated OCI-AML3 (iOCI3)-stimulation, 10,000 iOCI3 per well were added weekly.

##### IFN-γ ELISpot Assay

Response analyses by IFN-γ ELISpot assays were performed as reported previously (2). Briefly, NK and T cells were exposed to 100nM VTP-50469 for 14 to 16 days. The immune cells were then co-cultured with OCI-AML3 target cells ($6x10^4$ cells per 96-Well) at indicated E:T ratios for 16-20h in IFN-γ antibody-coated Multiscreen HTS plates (Merck-Millipore, Darmstadt, Germany). Immune cells cultivated without target cells served as negative controls. All reactions were set up in triplicates. After 20h, cells were removed and the tests developed as previously described (2). Plates were scanned and analyzed by ImmunoSpot Analyzer S5 Versa with ImmunoSpot software 7.0.15.1 (CTL Europe, Bonn, Germany).

##### Surface marker assessment by flow cytometry

Cells treated with drug or vehicle were centrifuged and washed twice with cold 1x phosphate buffered saline (PBS). Antibody staining was performed for 30min at 4°C in the dark. Stained cells were washed once, centrifuged at 1500 rpm for 5 minutes and resuspended in PBS supplemented with 2,2mM EDTA and DAPI (1µg/mL). Unstained cells served as negative control. Flow cytometry was performed on a FACS Canto II Flow Cytometer (BD Biosciences) or FACS Celesta (BD Biosciences).

##### Antibodies

| Antigen | Fluorochrome | Isotype | Clone | Dilution | Vendor |
|---|---|---|---|---|---|
| **BIOTIN** | PE | Recombinant human IgG1 | REA746 | 1:100 | Miltenyi Biotech |
| **CD11B** | APC | Mouse IgG1, κ | ICRF44 | 1:50 | BioLegend |
| **CD14** | BV711 | Mouse IgG2a, κ | M5E2 | 1:80 | Biolegend |
| **CD16** | FITC | Recombinant human IgG1 | REA423 | 1:500 | Miltenyi Biotec |
| **CD16** | BUV395 | Mouse IgG1, κ | 3G8 | 1:50 | BD Bioscience |
| **CD16** | PE | Mouse IgG1, κ | 3G8 | 1:80 | Biolegend |
| **CD19** | BB515 | Mouse IgG1, κ | HIB19 | 1:80 | BD Bioscience |
| **CD28** | FITC | Recombinant human IgG1 | REA612 | 1:50 | Miltenyi Biotec |
| **CD3** | APC | Mouse IgG2a, κ | HIT3A | 1:500 | Biolegend |
| **CD3** | PerCP | Mouse IgG1, κ | UCHT-1 | 1:50 | Biolegend |
| **CD3** | BUV395 | Mouse IgG1, κ | SK7 | 1:40 | BD Bioscience |
| **CD4** | FITC | Mouse IgG1, κ | RPA-T4 | 1:50 | BD |
| **CD4** | BV711 | Mouse IgG1, κ | SK3 | 1:40 | BD Bioscience |

| | | | | | |
|---|---|---|---|---|---|
| **CD45** | BV510 | Mouse IgG1, κ | HI30 | 1:40 | BD Bioscience |
| **CD45RA** | APC | Mouse IgG2b, κ | HI100 | 1:50 | BioLegend |
| **CD56** | APC | Mouse IgG1, κ | HCD56 | 1:50 | BioLegend |
| **CD56** | BV421 | Mouse IgG2b, κ | NCAM16.2 | 1:160 | BD Bioscience |
| **CD57** | PE | Recombinant human IgG1 | REA769 | 1:500 | Miltenyi Biotec |
| **CD62L** | PE | Recombinant human IgG1 | REA615 | 1:500 | Miltenyi Biotec |
| **CD69** | FITC | Mouse IgG1, κ | FN50 | 1:50 | BioLegend |
| **CD8** | FITC | Mouse IgG1, κ | HIT8a | 1:50 | BD |
| **CD8** | BUV737 | Mouse IgG1, κ | SK1 | 1:160 | BD Bioscience |
| **CD8** | APC | Mouse IgG1, κ | HIT8a | 1:50 | BD |
| **CLEC12A** | FITC | Recombinant human IgG1 | REA431 | 1:50 | MILTENYI BIOTEC |
| **CLEC12A** | APC | Recombinant human IgG1 | REA431 | 1:50 | Miltenyi Biotec |
| **CLEC12A** | BV605 | Mouse IgG2a, κ | 50C1 | 1:160 | BD Bioscience |
| **ISOTYPE CONTROL** | BV605 | Mouse IgG2a, κ | G155-178 | 1:160 | BD Bioscience |
| **KLRG1** | PE | Mouse IgG1, κ | Z7-205.rMAb | 1:50 | BD |
| **LAG3** | APC | Mouse IgG1, κ | 7H2C65 | 1:50 | BioLegend |
| **myc** | Biotin | Mouse IgG1 | SH1-26E7.1.3 | 1:100 | Miltenyi Biotech |
| **NKG2D** | APC | Mouse IgG1, κ | 1D11 | 1:50 | BioLegend |
| **PD-1** | APC | Mouse IgG1, κ | EH12.2H7 | 1:500 | BioLegend |
| **TIGIT** | APC | Recombinant human IgG1 | REA1004 | 1:50 | Miltenyi Biotec |
| **TIM3** | PE | Mouse IgG1, κ | 7D3 | 1:50 | BD |
| **TIM3** | PE | Recombinant human IgG1 | REA635 | 1:50 | Miltenyi Biotec |

**Morphological analysis**

For analysis of cell morphology following treatment with VTP-50469 $1x10^5$ AML or immune cells were harvested in 100µL PBS and cytospins (500rpm, 5 minutes) were performed in a Shandon Cytospin 4 (Thermo Fisher). After Giemsa staining according to standard procedures, images were taken at 40X (ATRA and menin inhibitor treatment, Supplemental-Figure 2) and 100X magnification (*in vivo* experiment, Figure 7).

**Chromatin Immunoprecipitation (ChIP)**

OCI-AML3 cells were treated for four days with 100nM VTP50469 or DMSO control. ChIP followed by DNA sequencing was performed as previously described (3). Briefly, cross-linking was performed with 1% formalin for 10min followed by cell lysis in SDS buffer. Sonication was used to fragment DNA. ChIP for KMT2A and Menin was performed using the anti-Menin antibody A300-105A (biomol) and anti-MLL1 A300-086A (biomol) specific to the respective modifications. Eluted DNA fragments were analyzed using DNA sequencing.

**Human Primary AML Blast Assay**

Primary AML samples were obtained from patients diagnosed at the University Medical Center, Mainz, or University Medical Center, Frankfurt, in accordance with the Declaration of Helsinki under Institutional Review-Board approved protocols and after written consent. The samples were selecte for *NPM1* mutations for the menin inhibitor treatment (wildtype served as controls). Primary cells M-01 to M-14 were isolated via Ficoll Density Centrifugation and cultured for six days in serum-free media supplemented with different cytokines (StemSpan Media (Stem Cell Technologies) + 1%P/S, 0.1% 2-Mercaptoetanol (Sigma), 100ng/mL SCF, 20ng/mL IL-3, 20ng/mL G-CSF, 50ng/mL FLT3L (PeproTech) in the presence of 100nM VTP-50469 or DMSO.. On day 6, CLEC12A expression was assessed by FC and the cells subjected to assessment in combination assays.

Primary AML samples F-01 and F-02, were cultivated in IMDM medium supplemented with ʟ-glutamine, 25mM HEPES, 1% P/S, a cytokine mix derived of 100 ng/mL human stem cell factor (hSCF), 20 ng/mL IL-3, 20 ng/mL IL-6 and 100 ng/ml FMS-like tyrosine kinase 3 ligand (FLT3L, all PeproTech) at a density of $2x10^6$ cells/mL in a 24-well plate (Thermo Fisher Scientific). One day post thawing, pAML cells were utilized for evaluation of CAR T cell functionality assays and flow cytometry. Percentage of AML blasts and CLEC12A surface expression were determined using flow cytometry with focus on the viable $CD45^{dim}$ cell population.

**Characteristics of primary AML patient cells**

| **ID** | **Molecular characteristics** | **Diagno-sis** | **Gen-der** | **AML blasts [%]** | **Age** | **Cyto-genetics** | **WBC/nl diagnosis** | **CLEC12A expression on D6 (DMSO-VTP)** |
|---|---|---|---|---|---|---|---|---|
| M-01 | *NPM1*$^{mut}$ *IDH2*$^{mut}$ | ND | m | 60-70% | 55 | 46, XY | 42,4 | 12,8%; 53,7% |
| M-02 | *NPM1*$^{mut}$ *FLT3*$^{ITD\ high}$ *IDH1*$^{mut}$ | ND | w | 99 | 23 | 46, XX | 227 | 13,3%; 30,6% |
| M-03 | *NPM1*$^{mut}$ *FLT3*$^{TKD}$ | ND | w | n.d. | 40 | 46, XX | 150 | 19,5%; 66% |
| M-04 | *NPM1*$^{mut}$ *FLT3*$^{TKD}$ | Re-lapse | m | >80% | 28 | 46, XY | 27,5 | 20,1%, 43,7% |
| M-05 | *NPM1*$^{mut}$ | ND | w | n.d. | 58 | 46, XX | 69,6 | 29,3%, 77% |
| M-06 | *NPM1*$^{mut}$ *FLT3*$^{ITD\ high}$ | ND | m | 90 | 70 | 46, XY | 67,4 | 50,8%; 83,8% |
| M-07 | *NPM1*$^{mut}$ *FLT3*$^{ITD\ high}$ | ND | w | 90 | 67 | 46, XX | 165 | 52,6%; 88,3% |
| M-08 | *NPM1*$^{mut}$ *FLT3*$^{ITD}$ | n.d. | n.d. | n.d. | n.d. | n.d. | n.d. | 59,1%, 60,8% |
| M-09 | *NPM1*$^{mut}$ | ND | w | 80% | 78 | 46, XX, (t4;11) (q31;21) | 86,6 | 68,6%, 91,4% |

| | | | | | | | | |
|---|---|---|---|---|---|---|---|---|
| M-10 | *NPM1*$^{mut}$ *FLT3*$^{ITD\ high}$ | ND | w | 80% | 65? | 46, XX | 37,3 | 71,9; 90,8% |
| M-11 | *NPM1*$^{mut}$*FLT3*$^{ITD\ low}$*IDH1*$^{mut}$ | ND | m | 80% | 59 | 46, XY | 0.6 | 73,2%; 83,8% |
| M-12 | *NPM1*$^{WT}$*FLT3*$^{ITD}$ | n.d. | n.d. | n.d. | n.d | n.d | n.d | 86,0 &, 98,4% |
| M-13 | inv16 | ND | m | 63% | 46 | 46, XY; inv(16) | 205 | 93,9%; 97,0% |
| M-14 | *IDH2*$^{mut}$ ASXL1$^{mut}$ RUNX1$^{mut}$ FLT3$^{mut}$ | ND | m | 92% | 55 | 47, XY; +11 | 151 | 4,9%, 5,8% |
| F-01 | *FLT3*$^{TKD}$ | ND | female | 85% | 28 | 46, XX | n.d. | 86% (baseline) |
| F-02 | none detected | ND | male | 86% | 54 | 46, XY | n.d. | 89% (baseline) |

**Design of the chimeric antigen receptor (CAR) targeting CLEC12A**

The CLEC12A-CAR sequence C12A.BBz was designed *in silico* and composed of a CD28 signal peptide (UniprotKB: P10747, amino acids 1-18), the variable light ($V_L$) and heavy chain ($V_H$) domain of the anti-CLEC12A scFv derived from Tepoditamab (MCLA-117, (4,5)) and linked by an $(G_4S)_3$-element, a Myc-tag sequence (EQKLISEEDL), a modified CD8α hinge domain as well as CD8α transmembrane domain (UniprotKB: P01732, amino acids 183-205), intracellular domains of 4-1BB (CD137, UniprotKB: Q07011, amino acids 214-255) and CD3ζ (UniprotKB: P20963, CD3ζ isoform 3, amino acids 52-163). Individual, codon-optimized CAR modules were *de novo* synthesized (TWIST Bioscience) and prepared for assembly using the Golden Gate cloning technique. All CAR modules were integrated into a modified version of the third-generation lentiviral transfer plasmid pSLCAR (Addgene #135991) (6) under control of a human EF-1α core promoter, resulting in the vector pSL-GG-C12A.BBz.

**Production of third-generation lentiviral vectors**

Production of VSV-G- and RD114TR-pseudotyped, self-inactivating lentiviral vectors (LV) was performed using a third-generation plasmid system and 25kDa, linear polyethyleneimine (PEI)-based (Polysciences, transfection grade) transfection of Lenti-X™ 293T cells (Takara Bio). Briefly, 293T cells were transfected at 70-80% confluence with the following plasmids: pSL-GG-C12A.BBz (transgene, 23µg), pcDNA3.g/p.4xCTE (Gag/Pol, 23µg), pRSV-REV (Rev, 11.5µg) and pCMVRDTR (RD114TR, 9.2µg) or pMD2.G (VSV-G, 3.5µg). Following harvest of virus-containing supernatant 40-46h after transduction, remaining cell debris was removed by centrifugation and filtration trough a 0.22µm filter system (Stericup® Quick Release Vacuum Filtration System, Merck Millipore) and viral vectors were concentrated in a sucrose cushion (20% w/v, in $PBS^{-/-}$) by centrifugation at 4500g and 4°C for 24h in an Avanti JXN-30 (Beckman Coulter). Concentrated lentiviral vectors were resuspended in NK MACS

(RD114TR) or RPMI 1640 (VSV-G) and stored at -80°C. Frozen virus batches were titrated using primary NK cells (RD114TR) or primary T cells (VSV-G).

**Transduction of primary immune cells**

Transduction with VSV-G-pseudotyped LV vectors was performed as recently described (7). In detail, primary T cells were seeded post isolation in 100µL TCMp inside a 96-well plate at a cell density of $1x10^6$ cells/mL and were activated by addition of 50 IU/mL IL-2 (PROLEUKIN S, Novartis Pharma, Basel, Switzerland) as well as human T-Activator CD3/CD28 DynaBeads (Gibco) at a ratio of 0.5:1 beads per cell. The following day, activated T cells were transduced with VSV-G-pseudotyped LV vectors (MOI = 1) diluted in RPMI 1640 by spin-infection at 800xg for 90 minutes at 32°C. Half of the medium was replaced 18-24h post transduction by fresh TCMp supplemented with 50 IU/mL IL-2. Subcultivation of the T cells to a cell density of $0.5x10^6$ cells/mL was performed every three to four days. Transduction of primary NK cells with RD114TR-pseudotyped LV vectors (MOI=1) was performed on day two post isolation as recently described (1)(8). CAR expression and phenotypes of engineered CAR T and CAR NK cells were monitored by FC (FACS Celesta & FACS Canto II, BD).

**Cytotoxicity of non-transduced or CAR NK/CAR T cells**

Functionality of generated CAR T and CAR NK cells as well as non-transduced immune cells was assessed in FC- and Firefly luciferase-based cytotoxicity assays against the AML cells lines OCI-AML2, OCI-AML3, MOLM13 and primary AML cells. For evaluation of menin inhibitor-mediated effects, target cells were pretreated with 100nM VTP-50469 (MedChem Express) or DMSO for six (primary AML blasts) or four days (AML cell lines) in respective media. AML cells were counted and an equal number of viable cells per well was plated for both conditions (menin inhibitor-pretreated and DMSO-pretreated) in a 96-well plate. Tumor cells were either luciferase-expressing (*FLUC*-OCI-AML3) or labeled with 50nM CFSE (Thermo Fisher Scientific) according to the manufacturer's instructions. Immune cells (+/- CAR) were added in indicated effector to target (E:T) ratios (maximum 10:1, lowest 1:27)). For all assays, AML cells cultivated without effector cells served as normalization control.

After 4 to 24h, CFSE-labeled cells were stained with 1µg/ml DAPI (AppliChem/Roth) and viable $DAPI^{low}$ cells counted using FC (FACS Celesta & FACS Canto II, BD Biosciences). Specific lysis was calculated normalizing the percentage of the dead target cells to the spontaneous lysis of target cells without effector cells (AML only) as described in formula S1.

$$specific\ lysis\ [\%] = \frac{(dead\ target\ cells\ [\%] - spontanous\ lysis\ [\%])}{(1 - \ spontanous\ lysis\ [\%])}$$

**Formula S1. Calculation of specific lysis in flow cytometry-based cytotoxicity assays.**

For the combination assays, viability and specific lysis was calculated as described in formula S2+3:

$$viability\ \ [\%] = \ 100x\ \frac{(viable\ cells)}{(viable\ AML\ cells\ DMSO\ only\ group)}$$

**Formula S2. Calculation of viability in flow cytometry-based combination assays.**

$$specific\ lysis\ [\%] = 100x\ (1 - \frac{(viable\ cells)}{(viable\ cells\ DMSO\ only\ group)})$$

**Formula S3 Calculation of specific lysis in flow cytometry-based combination assays.**

For *FLUC*- OCI-AML3 cells, 25µg/mL D-Luciferin (Revvity) was added with the effector cells. After 6 or 19h, absolute luminescence was quantified by an OMEGA plate reader (FluOstar OMEGA 0415, 2007) correlating to the luciferase activity of viable cells. Calculation of viability and lysis for luminescence-based assays was performed as described in formula S4+ S5.

$$viability\ [\%] = 100x\ \frac{(luminescence\ - luminescence\ background)}{(luminescence\ AML\ cell\ only\ control\ group - luminescence\ background)}$$

**Formula S4. Calculation of viability in luciferase-based cytotoxicity assays.**

$$lysis[\%] = 100x\ (1 - \frac{(luminescence\ - luminescence\ background)}{(luminescence\ AML\ cell\ only\ control\ group - luminescence\ background)})$$

**Formula S5. Calculation of specific lysis in luciferase-based cytotoxicity assays**

**Purity and phenotype of CAR T and -NK cells**

Purity of isolated NK cells was confirmed on the day of isolation by flow cytometry using a FACS Celesta (BD Biosciences). Briefly, 1-2x$10^5$ cells were washed with PBS and stained for 30 minutes at 4°C using the following antibody panel: anti-CD3-BUV395 (BD Bioscience), anti-CD14-BV711 (Biolegend), anti-CD16-PE (Biolegend), anti-CD19-BB515 (BD Biosciences), anti-CD45-BV510 (BD Biosciences) and anti-CD56-BV421 (BD Biosciences). Afterwards, cells were washed and stained using 7-AAD (BD Pharmingen) for discrimination of dead and viable cells.

CAR expression and phenotype of primary T and NK cells were monitored by flow cytometry using a FACS Celesta and FACS Canto II, following the staining protocols described above. Characterization was performed using the following antibodies: anti-Myc-Biotin (Miltenyi Biotech) and anti-Biotin-PE (Miltenyi Biotech) for CAR detection; anti-CD3-PerCP (Biolegend), anti-CD4-BV711 (BD Bioscience) and anti-CD8-BUV737 (BD Bioscience) for T cell subtypes; anti-CD16-BUV395 (BD Bioscience) and anti-CD56-BV711 (BD BioscienceHorizon) for NK cell subtypes. All data was analyzed using FlowJo Version 10 software.

**Single Cell mRNA Sequencing**

Viability analysis, single cell capturing and mRNA isolation was performed with a BD Rhapsody Single Cell analysis system (BD Biosciences) following manufacturer‘s guidelines (Single Cell Capture and cDNA Synthesis with the BD Rhapsody™ Single-Cell Analysis System). Each sample was tagged with a unique sample tag allowing to pool samples from different donors on the same single cell cartridge (1 cartridge T cells, 1 cartridge NK cells) using the BD™ Human Sample Multiplexing Kit (BD Biosciences, Catalog No: 633781) according to the manufacturer’s protocol. DNA libraries for Whole Transcriptome Analysis (WTA) and Sample Tags were constructed according to the BD Rhapsody System mRNA Whole Transcriptome analysis (WTA) and Sample Tag Library Preparation Protocol with the BD WTA Amplification Kit (BD Biosciences, Franklin Lakes, USA, Catalog No.: 633801). The sample preparation was performed in cooperation with the Research Center for Immunotherapy (FZI) Core Facility NGS of the Johannes Gutenberg-University Mainz. Sequencing was performed by Novogene Co. Ltd. (Cambridge, UK) of a minimum of 2,000 cells per sample at each 30,000 reads.

**Single Cell Sequencing Data Analysis**

The initial data processing, quality control, and integration followed the data analysis protocol described by Nedwed et al.(9). Short read sequences were quantified using the BD Rhapsody analysis pipeline and the dataset was annotated to gene-level information based on ENSEMBL release 112 for *Homo sapiens*. Quality control was performed on each dataset independently to remove poor-quality cells with scater v1.35.0(10). The proportion of mitochondrial gene content was used as a proxy for damaged cells, using three median absolute deviations as a threshold, following the recommendations of the OSCA resource (https://bioconductor.org/books/release/OSCA/)(11). Doublet detection was performed using scDblFinder v1.21.0 (12). Normalization of cell-specific biases was performed on the sets of cells passing the quality control filters using the deconvolution method of Lun et al.(13). Counts were divided by size factors to obtain normalized expression values that were log-transformed after adding a pseudocount of one. Integration of different biological samples was conducted using batchelor v1.23.0, specifically using the MNN method (14). Highly variable genes were identified on the pooled set of cells after decomposing the per-gene variability into technical and biological components based on a fitted mean-variance trend. Next, we performed dimension reduction and clustering. Principal Component Analysis (PCA) was performed, and provided as initialization to the t-SNE algorithm (15) to obtain a reduced dimensionality representation of the data. Clustering was performed using the 13094 (NK cells dataset) / 16415 (T cells dataset) most highly variable genes (HVGs), building a shared nearest neighbor graph (16). The Louvain community finding algorithm (19) was applied to determine cluster memberships. Initial cell type annotation was performed automatically using SingleR v2.9.0 (17) with the MonacoImmuneData reference dataset from celldex v1.17.0 (17). This annotation was further manually refined using well-established cell type marker genes from the literature (8,18–23), with visualization and adjustments carried out using iSEE v2.19.0 and iSEEfier v1.5.0 (24),(25). Complementary exploration was conducted using iSEE v2.19.0, which was adopted to generate most single-cell data visualizations included. Pseudobulk differential expression analysis was performed with muscat v1.21.0 (26), using the default model for modeling the counts. Multiple testing correction was applied using Benjamini-Hochberg (BH) correction, and genes were considered differentially expressed (DEGs) for p-value <0.05. Functional enrichment analysis was conducted using mosdef v1.3.0 (27), implementing topGO (28) for Gene Ontology enrichment analysis. The "elim" algorithm was used and the enrichment was conducted for the Biological Process ontology, with differentially expressed genes as input and all detected genes as the background set. Differential Abundance Analysis was performed with speckle v1.7.0 (29), comparing cell proportions across the treatments, with significance determined at FDR < 0.05.

**Data Analysis and Statistical Methods**

Statistics were calculated with Prism, Version 10 software (GraphPad Prism). Two-sided student t-tests were used to assess significance in the *in vitro* experiments and the leukemia burden *in vivo* experiment. For *in vitro* experiments, cytotoxicity and proliferation assays were performed in technical triplicates, immunophenotyping in technical duplicates. The means of the individual experiments were obtained and used for the calculation of significances between treatment groups. Kaplan-Meier Log-Rank was used for survival analysis. P values <0.05 were considered as significant with *, P<0.05; ** P< 0.01, *** p<0.001.

**References Supplement**


1. Reindl LM, Jalili L, Bexte T, Harenkamp S, Thul S, Hehlgans S, et al. Precision targeting of rhabdomyosarcoma by combining primary CAR NK cells and radiotherapy. J Immunother Cancer. 2025;13:e011330.

2. Lennerz V, Fatho M, Gentilini C, Frye RA, Lifke A, Ferel D, et al. The response of autologous T cells to a human melanoma is dominated by mutated neoantigens. Proc Natl Acad Sci U S A. 2005;102:16013–8.

3. Kühn MWM, Song E, Feng Z, Sinha A, Chen C-W, Deshpande AJ, et al. Targeting Chromatin Regulators Inhibits Leukemogenic Gene Expression in NPM1 Mutant Leukemia. Cancer Discov.; 2016;6:1166–81.

4. Raybould MIJ, Marks C, Lewis AP, Shi J, Bujotzek A, Taddese B, et al. Thera-SAbDab: the Therapeutic Structural Antibody Database. Nucleic Acids Res. 2020;48:D383–8.

5. van Loo PF, Hangalapura BN, Thordardottir S, Gibbins JD, Veninga H, Hendriks LJA, et al. MCLA-117, a CLEC12AxCD3 bispecific antibody targeting a leukaemic stem cell antigen, induces T cell-mediated AML blast lysis. Expert Opin Biol Ther. 2019;19:721–33.

6. Bloemberg D, Nguyen T, MacLean S, Zafer A, Gadoury C, Gurnani K, et al. A High-Throughput Method for Characterizing Novel Chimeric Antigen Receptors in Jurkat Cells. Mol Ther Methods Clin Dev. 2020;16:238–54.

7. Prommersberger S, Hudecek M, Nerreter T. Antibody-Based CAR T Cells Produced by Lentiviral Transduction. Curr Protoc Immunol. 2020;128:e93.

8. Bexte T, Albinger N, Al Ajami A, Wendel P, Buchinger L, Gessner A, et al. CRISPR/Cas9 editing of NKG2A improves the efficacy of primary CD33-directed chimeric antigen receptor natural killer cells. Nat Commun. 2024;15:8439.

9. Nedwed AS, Helbich SS, Braband KL, Volkmar M, Delacher M, Marini F. Using combined single-cell gene expression, TCR sequencing and cell surface protein barcoding to characterize and track CD4+ T cell clones from murine tissues. Front Immunol [Internet]. Frontiers; 2023 [cited 2025 Aug 14];14.

10. McCarthy DJ, Campbell KR, Lun ATL, Wills QF. Scater: pre-processing, quality control, normalization and visualization of single-cell RNA-seq data in R. Bioinformatics. 2017;33:1179–86.

11. Amezquita RA, Lun ATL, Becht E, Carey VJ, Carpp LN, Geistlinger L, et al. Orchestrating single-cell analysis with Bioconductor. Nat Methods; 2020;17:137–45.

12. Germain P-L, Lun A, Meixide CG, Macnair W, Robinson MD. Doublet identification in single-cell sequencing data using *scDblFinder* [Internet]. F1000Research; 2022 [cited 2025 Aug 14].

13. L. Lun AT, Bach K, Marioni JC. Pooling across cells to normalize single-cell RNA sequencing data with many zero counts. Genome Biology. 2016;17:75.

14. Haghverdi L, Lun ATL, Morgan MD, Marioni JC. Batch effects in single-cell RNA-sequencing data are corrected by matching mutual nearest neighbors. Nat Biotechnol. 2018;36:421–7.

15. Maaten L van der, Hinton G. Visualizing Data using t-SNE. Journal of Machine Learning Research. 2008;9:2579–605.

16. Xu C, Su Z. Identification of cell types from single-cell transcriptomes using a novel clustering method. Bioinformatics. 2015;31:1974–80.

17. Aran D, Looney AP, Liu L, Wu E, Fong V, Hsu A, et al. Reference-based analysis of lung single-cell sequencing reveals a transitional profibrotic macrophage. Nat Immunol. 2019;20:163–72.

18. Galletti G, De Simone G, Mazza EMC, Puccio S, Mezzanotte C, Bi TM, et al. Two subsets of stem-like CD8+ memory T cell progenitors with distinct fate commitments in humans. Nat Immunol. 2020;21:1552–62.

19. Rebuffet L, Melsen JE, Escalière B, Basurto-Lozada D, Bhandoola A, Björkström NK, et al. High-dimensional single-cell analysis of human natural killer cell heterogeneity. Nat Immunol. 2024;25:1474–88.

20. Mullan KA, de Vrij N, Valkiers S, Meysman P. Current annotation strategies for T cell phenotyping of single-cell RNA-seq data. Front Immunol. 2023;14:1306169.

21. Terekhova M, Swain A, Bohacova P, Aladyeva E, Arthur L, Laha A, et al. Single-cell atlas of healthy human blood unveils age-related loss of NKG2C+GZMB−CD8+ memory T cells and accumulation of type 2 memory T cells. Immunity. 2023;56:2836-2854.e9.

22. Buggert M, Price DA, Mackay LK, Betts MR. Human circulating and tissue-resident memory CD8+ T cells. Nat Immunol. 2023;24:1076–86.

23. Chu Y, Dai E, Li Y, Han G, Pei G, Ingram DR, et al. Pan-cancer T cell atlas links a cellular stress response state to immunotherapy resistance. Nat Med. 2023;29:1550–62.

24. Rue-Albrecht K, Marini F, Soneson C, Lun ATL. iSEE: Interactive SummarizedExperiment Explorer [Internet]. F1000Research; 2018 [cited 2025 Aug 14]. Available from: https://f1000research.com/articles/7-741

25. iSEEfier [Internet]. Bioconductor. [cited 2025 Aug 14]. Available from: http://bioconductor.org/packages/iSEEfier/

26. Crowell HL, Soneson C, Germain P-L, Calini D, Collin L, Raposo C, et al. muscat detects subpopulation-specific state transitions from multi-sample multi-condition single-cell transcriptomics data. Nat Commun. 2020;11:6077.

27. mosdef [Internet]. Bioconductor. [cited 2025 Aug 14]. Available from: http://bioconductor.org/packages/mosdef/

28. Alexa A, Rahnenführer J, Lengauer T. Improved scoring of functional groups from gene expression data by decorrelating GO graph structure. Bioinformatics. 2006;22:1600–7.

29. Phipson B, Sim CB, Porrello ER, Hewitt AW, Powell J, Oshlack A. propeller: testing for differences in cell type proportions in single cell data. Bioinformatics. 2022;38:4720–6.

30. Krivtsov AV, Evans K, Gadrey JY, Eschle BK, Hatton C, Uckelmann HJ, et al. A Menin-MLL Inhibitor Induces Specific Chromatin Changes and Eradicates Disease in Models of MLL-Rearranged Leukemia. Cancer Cell. 2019;36:660-673.e11.

#### Supplemental Figures

**Supplemental-Figure 1:**

**A Induction of CLEC12A surface expression on day 4.** *KMT2A*-r, *NPM1*$^{mut}$ and wildtype cell lines were treated with 100nM VTP-50469 for 4 days. CLEC12A surface expression was determined by FC in viable (DAPI$^{low}$) cells. Displayed is the mean of each experiment, all performed in technical triplicates. Significance by unpaired t test, ** p<0.01, *** p<0.001. **B Induction of CLEC12A surface expression on day 7 displayed as MFI and fold change.** *KMT2A*-r, *NPM1*$^{mut}$ and wildtype cell lines were treated with 100nM VTP-50469 for 7 days. CLEC12A surface expression was determined by FC in viable (DAPI$^{low}$) cells. Displayed is the mean flow intensity (MFI) or the fold change of MFI of menin-i versus DMSO. Each symbol represents the mean of one individual experiment, all performed in technical triplicates. Significance by unpaired t-test, ** p<0.01, *** p<0.001. **C High CLEC12A surface expression is sustained in *NPM1*$^{mut}$ cells with adaptive menin inhibitor resistance**. Non-genetic menin inhibitor resistance was developed by increasing the MI-503 concentration in three independent single-cell OCI-AML3 clones over 5 months. CLEC12A expression was assessed by FC and depicts a sustained high CLEC12A expression compared to native OCI-AML3 counterparts. Significance by unpaired t-test, ** p<0.01, *** p< 0.001. **D ChIP sequencing upon menin-inhibition detects no binding of Menin or KMT2A on the locus.** Chromatin Immunoprecipitation (ChIP) Sequencing using anti-Menin or anti-MLL antibodies of OCI-AML3 treated with 100nM VTP-50469 for four days. To the left Menin and MLL binding at the *MEIS1* locus as control, to the right Menin and MLL binding at the *CLEC12A* binding site. **E ChIP Sequencing upon menin-inhibition detects no binding of Menin-or KMT2A on the locus in *KMT2A-r* MOLM13.** Re-analysis of previously published, publicly available data (30). To the left Menin and MLL binding at the *MEIS1* locus as control, to the right Menin and MLL binding at the *CLEC12A* binding site.

**Supplemental-Figure 2:**

**A+B No induction of CLEC12A surface expression upon ATRA treatment.** *KMT2A*-r, *NPM1*$^{mut}$ and wildtype cell lines were treated with 50nM ATRA for 7 days. CLEC12A or CD11b surface expression was assessed by FC in viable (DAPI$^{low}$) cells. Displayed is the mean of each experiment, all performed in technical triplicates. Significance by unpaired t-test, *** p<0.001. **C Cytomorphology depicts differences in differentiation upon VTP50469 or ATRA treatment.** *KMT2A*-r, *NPM1*$^{mut}$ and wildtype cell lines were treated with 100nM VTP-50469 or 50nM ATRA for 7 days. Cytomorphology was evaluated at 40X magnification.

**Supplemental-Figure 3:**

**A Effects of menin-inhibition on the proliferation of T cells stimulated with irradiated OCI-AML3.** T cells were stimulated with irradiated OCI-AML3 and 50IU/ml IL-2 as indicated in Figure 2A. Treatment with different concentrations of VTP-50469 (100nM max, 1:2 dilutions) for a total of 21 days. Displayed is the FC- readout of viable cells (DAPI$^{low}$) on day 7, 14 and 21 in comparison to DMSO. OCI-AML3 were treated with the same concentrations of the menin inhibitor as control. Summary of three experiments, each performed in technical triplicates, displayed are the mean+SEM for each concentration.

**B Menin-inhibition does not affect the phenotype of T cells.** T cells treatment as depicted in Figure 2A. Surface expression of the immune checkpoints PD-1, KLRG-1, TIM-3, TIGIT and LAG-3 on viable ($DAPI^{low}$) cells was assessed by FC on day 7 and 14, as well as of CD45RA, CD62L, CD57 and CD28 on day 7, 14, and 21. Each symbol represents one biological donor in technical duplicates. Statistical analysis by unpaired t-test.

**Supplemental-Figure 4:**

**A Menin-inhibition does not affect the phenotype of NK cells.** Surface expression of the immune checkpoints PD-1, KLRG-1, TIM-3, TIGIT and LAG-3 on viable ($DAPI^{low}$) NK cells on day 7 and 14, as well as of NKG2A, CD16, CD57 and CD69 on day 7, 14, and 21 by flow cytometry. CD56 is displayed as MFI. Each symbol represents one biological donor, measured in technical duplicates. Statistical analysis by unpaired t-test. **B Cytomorphology of T or NK cells upon menin-inhibition.** Cytomorphology after 21 days of menin-inhibition at 40X magnification.

**Supplemental-Figure 5:**

**A Menin-inhibition has no influence on the IFN-γ response of T cells.** T cells were exposed to 100nM VTP-50469 for 13 or 14 days, respectively. Cells were stimulated with viable OCI-AML3 and the IFN-γ release detected by ELISpot assays after 20h. Effector cells without targets served as negative control. **B Reactive cell count of T cells.** Reactive T cell spots were counted using ImmunoSpot software. Significance by t-test. **C NK cells exhibit an intact IFN-γ response upon menin-inhibition.** NK cells were exposed to 100nM or 187nM VTP-50469 for 15 or 16 days, respectively. Cells were stimulated with viable OCI-AML3 and the IFN-γ release detected by ELISpot assays. Effector cells without targets served as negative control. **D Reactive cell count of NK cells.** Reactive NK cell spots were counted using ImmunoSpot software. Significance by t-test. **E Setup of the cytotoxicity assay for NT T and NK cells.** NT T and NK cells were cultivated as indicated in Figure 2A and exposed to 100nM VTP-50469 for 7 days. NT T cells were simultaneously primed with lethally irradiated OCI-AML3 cells for 7 days to create alloreactivity. *FLUC*-OCI-AML3 cells were plated in a constant number per well and exposed to T or NK cells in different E:T- ratios. D-Luciferin was added as substrate for luminescence. After 19 hours, cytotoxicity was determined as the luminescence of the indicated group relative to the luminescence of *FLUC*-OCI-AML3 cells only as control. Figure made in BioRender.com.

**Supplemental-Figure 6:**

**A Marker panel defining the T cell subtypes.** Expression of various immune cell markers across the 17 T cell subpopulations. **B MDS-Plot.** Multidimensional-Scaling (MDS) plot visualizing the similarity or dissimilarity between different T cell samples. Samples with similar expression patterns cluster together while dissimilar samples separate. **C t-SNE split by donor.** t-SNE plot of T cells displaying the different T cell subclusters, split for the four different donors. **D No significant changes in differential gene expression**. Volcano plots of the differential gene expression of the most exhausted ($CD8^+$ effector cells, exhaustion) and most naïve ($CD4^+$ stem cell memory cells) T cell subtypes were exemplarily chosen to demonstrate that there were no differentially expressed genes. The only significantly altered gene was in the $CD8^+$ pre-T effector cell cluster (*LINC02446*).

**Supplemental-Figure 7:**

**A Differential expression of naïve T cell markers.** Display of selected naïve markers in the $CD4^+$ stem cell memory cluster (most naïve T cell subtype identified), split both by donor and split by condition. No significant differences were found between menin inhibitor- or DMSO-treated cells. **B Differential expression of cytotoxic markers.** Display of selected cytotoxic markers in the $CD8^+$ T cells with exhaustion cluster (most cytotoxic T cell subtype identified), split both by donor and split by condition. No significant differences were found between menin inhibitor- or DMSO-treated cells. **C Differential expression of exhaustion markers.** Display of selected exhaustion markers in the $CD8^+$ T cells with exhaustion cluster (most exhausted T cell subtype identified), split both by donor and split by condition. No significant differences were found between menin inhibitor- or DMSO-treated cells.

**Supplemental-Figure 8:**

**A Marker panel defining the NK cell subtypes.** Expression of various immune cell markers across the 5 NK cell subpopulations as previously described (19). **B MDS-Plot.** Multidimensional-Scaling (MDS) plot visualizing the similarity or dissimilarity between NK cell samples. Samples with similar expression patterns cluster together while dissimilar samples separate. **C t-SNE split by donor.** t-SNE plot of NK cells displaying the different NK cell subclusters, split for the four different donors. **D Differential expression of selected genes for NK cell subtypes.** Display of selected markers defining a cytotoxic or migration signature in NK cells, split both by donor and split by condition. No significant differences were found between menin inhibitor- or DMSO-treated cells.

**Supplemental-Figure 9:**

**Differential expression of typical NK cell receptor genes for the five NK cell subtypes identified.** Display of selected genes encoding typical NK cell receptors, split both by donor and split by condition. No significant differences were found between menin inhibitor- or DMSO-treated cells.

**Supplemental-Figure 10**

**A CAR T cells show strong cytotoxicity against OCI-AML2 cells.** CLEC12A-CAR T cell functionality was investigated against OCI-AML2 following 24h co-culture at increasing E:T-ratios 0.5:1 to 5:1 (n=5-8). **B Gating strategy for characterization of primary AML.** Primary AML blasts were identified based on size (FSC) and CD45 expression level. **C CLEC12A expression of primary AML cells.** CLEC12A expression was assessed by FC for all viable cells based on single cell gate, including lymphocytes and blast cells.

**Supplemental-Figure 11**

**A CLEC12A expression of *FLUC* OCI-AML3 is increased after 4, 5 and 7 days.** Cells were exposed to 100nM VTP-50469 or DMSO for 4, 5 or 7 days. CLEC12A surface expression was determined by measuring viable ($DAPI^{low}$) cells by FC. Summary of three experiments in technical triplicates, significance by unpaired t-test, *** $p<0.001$. **B Summary of 5 donors evaluating the combination of menin-inhibition with CAR T-cell therapy.** *FLUC*-OCI-AML3 were treated with DMSO (4days+19h),

VTP-50469 100nM (4days+19h), CAR T (19h) or menin inhibitor (4 days+19h) + CAR T cells (19h) as indicated in Figure 5A. Viability was determined by luminescence in comparison to DMSO. Displayed is the summary of 5 donors for the E:T-ratios 1:1 and 1:3. Each symbol represents one experiment with a different biological immune cell donor, each experiment was performed in technical triplicates. AML cell viability is displayed, with killing defined as 100% − viability. Analysis by unpaired t-test, * p<0.05, ** p<0.01, *** p<0.001. **C CAR T cells exhibit a superior specific lysis against menin inhibitor pre-treated AML cells.** Luminescence was normalized to either DMSO- or VTP-50469 pre-treated OCI-AML3 cells without added CAR cells (control) to exclude the anti-leukemic effects of the menin inhibitor alone. Summary of five experiments in each technical triplicates. Each symbol reflects one different biological CAR T cell donor. Significance by unpaired t-test, * p<0.05. **D Summary of 5 donors evaluating the combination of menin-inhibition with NT T-cell therapy.** *FLUC*-OCI-AML3 were treated with DMSO (4days+19h), VTP-50469 100nM (4days+19h), NT T (19h) or menin inhibitor (4days+19h) + NT T cells (19h) as indicated in Figure 5A. Viability was determined by luminescence in comparison to the DMSO control. Displayed is the summary of 5 donors for the E:T-ratios 1:1 and 1:3. Each symbol represents one donor, each experiment was performed in technical triplicates. AML cell viability is displayed, with killing defined as 100% − viability. Analysis by unpaired t-test, * p<0.05, ** p<0.01, *** p<0.001. **E NT T cells show no specific lysis nor differences between menin inhibitor or DMSO pre-treatment.** Luminescence signal from the specific groups normalized either to the DMSO- or VTP-50469 pre-treated OCI-AML3 cells (control group) to exclude the anti-leukemic effects of the menin inhibitor alone. Summary of five experiments in each technical triplicates. Each symbol reflects one different biological CAR T cell donor. Significance by unpaired t-test, * p<0.05. **F The combination of menin-inhibition and CLEC12A-CAR T cells exhibits superior anti-leukemic effects after 6 hours of co-culture time.** Set-up of the experiment as described in A with readout after 6 hours. OCI-AML3 cells were stained with CSFE and the number of CSFE$^{high}$ cells counted by FC and normalized to the DMSO-pretreated OCI-AML3 only group. Display of three individual donors for the E:T-ratio 3:1, each experiment in technical triplicates. Significance by t-test, * p<0.05 ** p<0.01, ** p<0.001. **G Specific lysis against DMSO- or menin inhibitor pre-treated AML cells after 6 hours.** The number of CSFE$^{high}$ OCI-AML3 cells of each group was normalized to either the DMSO- or VTP-50469 pretreated OCI-AML3 only group (control) to exclude anti-leukemic effects of the menin inhibitor alone. Summary of three experiments in each technical triplicates. Each symbol reflects one CAR T cell donor. Significance by unpaired t-test.

**Supplemental-Figure 12**

**A Summary of four donors evaluating the combination of menin-inhibition with CAR T-cell therapy in *KMT2A*-r MOLM13 cells.** MOLM13 cells were pretreated with DMSO or VTP-50469 100nM for four days. On day four MOLM13 were stained with CSFE and CAR T cells were added to the respective conditions as indicated in Figure 5A. Readout was after coculture for 6h or 19h. Viability was determined detecting the CSFE$^{high}$ MOLM13 cells of respective groups relative to the DMSO pretreated MOLM13 only group. Displayed is the summary of four donors for the E:T-ratios 1:3 and 1:9. Each symbol represents one biological T cell donor, each independent experiment was performed in technical triplicates. Analysis by unpaired t test, * p<0.05, ** p<0.01, *** p<0.001. **B Anti-leukemic activity of**

**combined menin-inhibition with CLEC12A-directed CAR T cells after 19h.** Display of the independent experiments summarized in A for the E:T-ratio of 1:3, each with a different biological T cell donor. Analysis by unpaired t test, * p<0.05, ** p<0.01, *** p<0.001. **C Specific lysis against DMSO- or menin inhibitor- pretreated MOLM13 cells after 19 hours.** The number of $CSFE^{high}$ MOLM13 cells of each group was normalized to either the DMSO- or VTP-50469 pretreated MOLM13 only group (control), respectively, to exclude the anti-leukemic effects of the menin inhibitor alone. Summary of five experiments in each technical triplicates. Each symbol reflects one CAR T cell donor. Significance by unpaired t-test. **D Combination of menin-inhibition and CAR T cells after 6h.** Display of the experiments from A for the individual donors in an E:T-ratio of 3:1. Statistical analysis by unpaired t test, * p<0.05, ** p<0.01, *** p<0.001. **E Specific lysis against DMSO- or menin inhibitor- pretreated MOLM13 cells after 6 hours.** The number of $CSFE^{high}$ MOLM13 cells of each group was normalized to either the DMSO- or VTP-50469 pretreated MOLM13 only group (control), respectively, to exclude anti-leukemic effects of the menin inhibitor alone. Summary of three experiments in each technical triplicates. Each symbol reflects one CAR T cell donor. Significance by unpaired t-test. **F Combination of menin-inhibition and NT T cells after 19h.** MOLM13 were treated with DMSO (4days+19h), VTP-50469 100nM (4days+19h), NT T (19h) or the combination of menin inhibitor (4days+19h) and NT T cells (19h) as indicated in Figure 5A. Viability was determined detecting the $CSFE^{high}$ MOLM13 of the respective groups in comparison to the DMSO- pretreated MOLM13 only group, displayed for the E:T-ratio of 1:3. Each experiment was performed in technical triplicates. Analysis by unpaired t-test, * p<0.05, ** p<0.01.

**Supplemental-Figure 13**

**A NT NK cells exhibit alloreactive activity with additive anti-leukemic effects in combination with menin-inhibition.** *FLUC*-OCI-AML3 cells were exposed to 100nM VTP50469 or DMSO as previously described and NT NK cells added as effector cells after 4 days. Displayed is the E:T-ratio of 1:1, both for the single treatment and the combination with menin-inhibition. Five independent experiments with different biological donors, each experiment in technical triplicates. Significance by unpaired t-test, *** p<0.001. **B Summary of 5 donors evaluating the combination of menin-inhibition with CLEC12A-directed CAR NK-cell therapy.** *FLUC*-OCI-AML3 were treated with DMSO (4days+19h), VTP-50469 100nM (4days+19h), CAR NK (19h) or menin inhibitor (4days+19h) + CAR NK cells (19h) as indicated in Figure 5A. Viability was determined by luminescence in comparison to DMSO. Displayed is the summary of 5 donors for the E:T-ratios 1:1 and 1:3. Each symbol represents one different biological immune cell donor, each experiment was performed in technical triplicates. Analysis by unpaired t- test, * p<0.05, ** p<0.01, *** p<0.001. **C CAR NK cells exhibit similar lysis to menin inhibitor pre-treated AML cells.** Luminescence was normalized to either DMSO- or VTP-50469- pretreated OCI-AML3 cells without added CAR cells (control), respectively, to exclude the anti-leukemic effects of the menin inhibitor alone. Summary of five experiments in each technical triplicates. Each symbol reflects one CAR NK cell donor. Significance by unpaired t-test, * p<0.05. **D Summary of 5 donors evaluating the combination of menin-inhibition with NT NK cells** *FLUC*-OCI-AML3 were treated with DMSO (4days19h), VTP-50469 100nM (4days19h), NT NK (19h) or menin inhibitor + NT NK cells (4days+19h) as indicated in Figure 5A. Viability was determined by luminescence normalized to DMSO. Displayed is the summary of 5 donors for the E:T-ratios 1:1 and 1:3. Each symbol represents one different biological immune cell donor, each experiment was performed in technical triplicates. Analysis by unpaired t- test, * p<0.05, **

p<0.01, *** p<0.001. **E NT NK cells exhibit similar lysis to menin inhibitor pre-treated AML cells.** Luminescence was normalized to either DMSO- or VTP-50469 pre-treated OCI-AML3 cells without added NT cells (control) to exclude the anti-leukemic effects of the menin inhibitor alone. Summary of five experiments in each technical triplicates. Each symbol reflects one different biological immune cell donor. Significance by unpaired t-test, * p<0.05. **F No differences between phenotype of CAR NK cells and NT NK cells and no effect of menin-inhibition on the phenotype of CAR NK after 7 days.** Comparison of CLEC12A, CD16, PD-1 and KLRG-1 surface expression of CAR NK and NT NK cells as well as of CD16, PD-1 and KLRG1 on CAR NK cells after 7 days of exposure to 100nM VTP-50469 or DMSO. Surface expression was assessed by FC in viable cells. Each symbol represents one donor in technical duplicates. Statistical analysis by unpaired t-test. **G No differences in surface expression between NT and CAR T cells upon menin-inhibition.** CLEC12A, CD4, CD8, PD-1 and KLRG-1 surface expression was assessed by FC on day 0 between NT and CAR T cells as well as after 7 days of treatment with 100nM VTP-50469 or DMSO in CAR and NT T cells by flow cytometry. Each symbol represents one donor in technical duplicates. Significance by unpaired t-test, * p<0.05.

**Supplemental-Figure 14**

**A *In vitro* combination assay using the CAR T cell donor from the *in vivo* experiment**. Experimental design as described in Figure 5A. MOLM13 cells were pretreated for 4 days with the menin inhibitor. The combined anti-leukemic activity was assessed 19h after the addition of the CLEC12A-CAR T cells measuring the CSFE$^{high}$ viable MOLM13 cells in respective groups relative to the MOLM13 only group. Statistical analysis by unpaired t test, *** p<0.001.

**B Gating strategy for the leukemic engraftment experiment.** Murine bone marrow cells were isolated and human CD45 and human CD3 staining performed after red blood cell lysis. Readout was FC-based. The analysis first detected viable cells, then excluded CD3$^+$ CAR T cells in respective groups. AML blasts were the remaining human CD45$^+$ cells. The engraftment percentage was calculated as human CD3$^-$ CD45$^+$ cells of all viable cells. **C Engraftment on day 10 of the survival experiments.** FC-analysis of viable human CD45$^+$ cells in the peripheral blood of day 10 of the survival experiments (day before the CAR T cell injection).

#### Supplemental-Figure 1

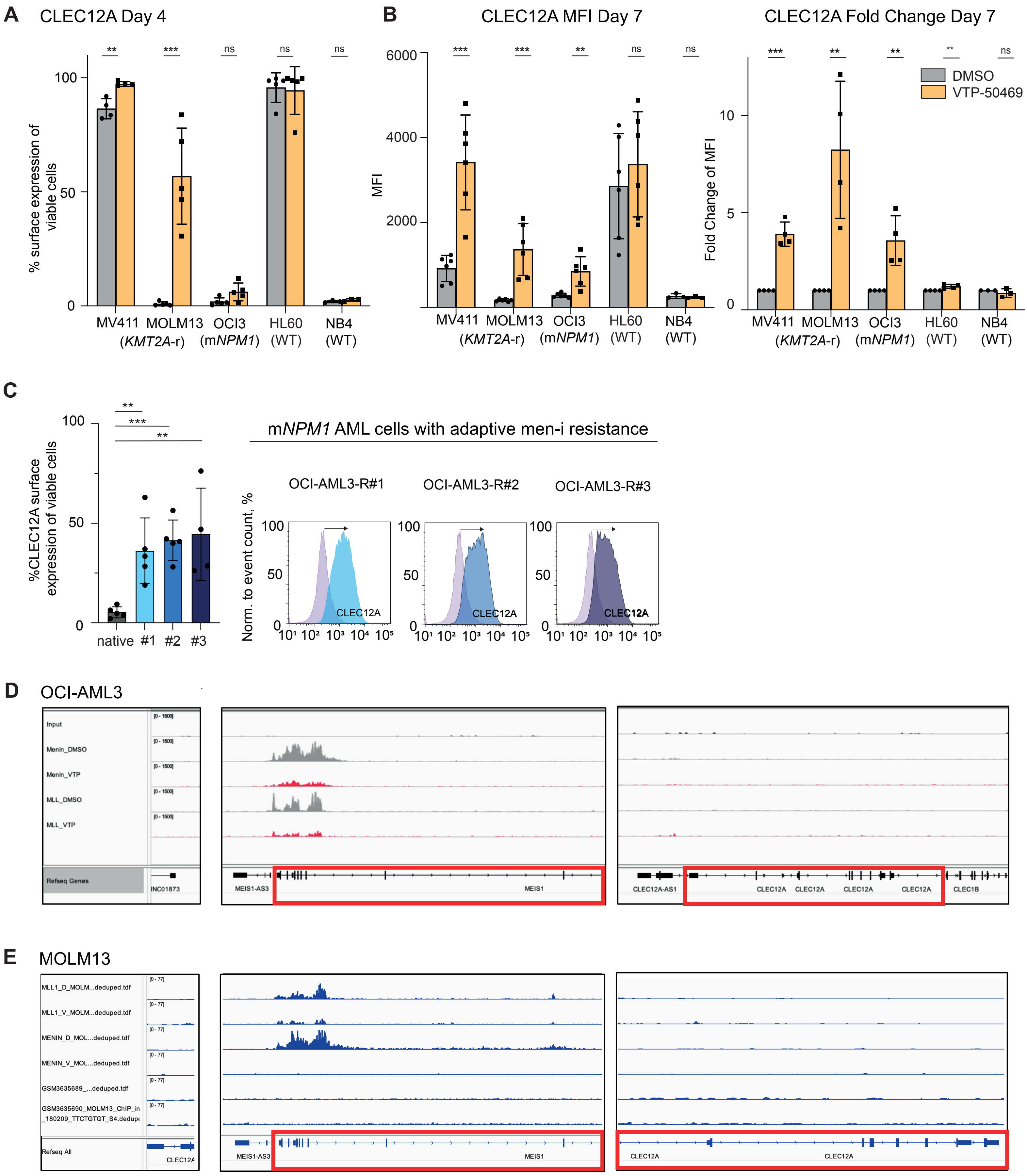

#### Supplemental-Figure 2

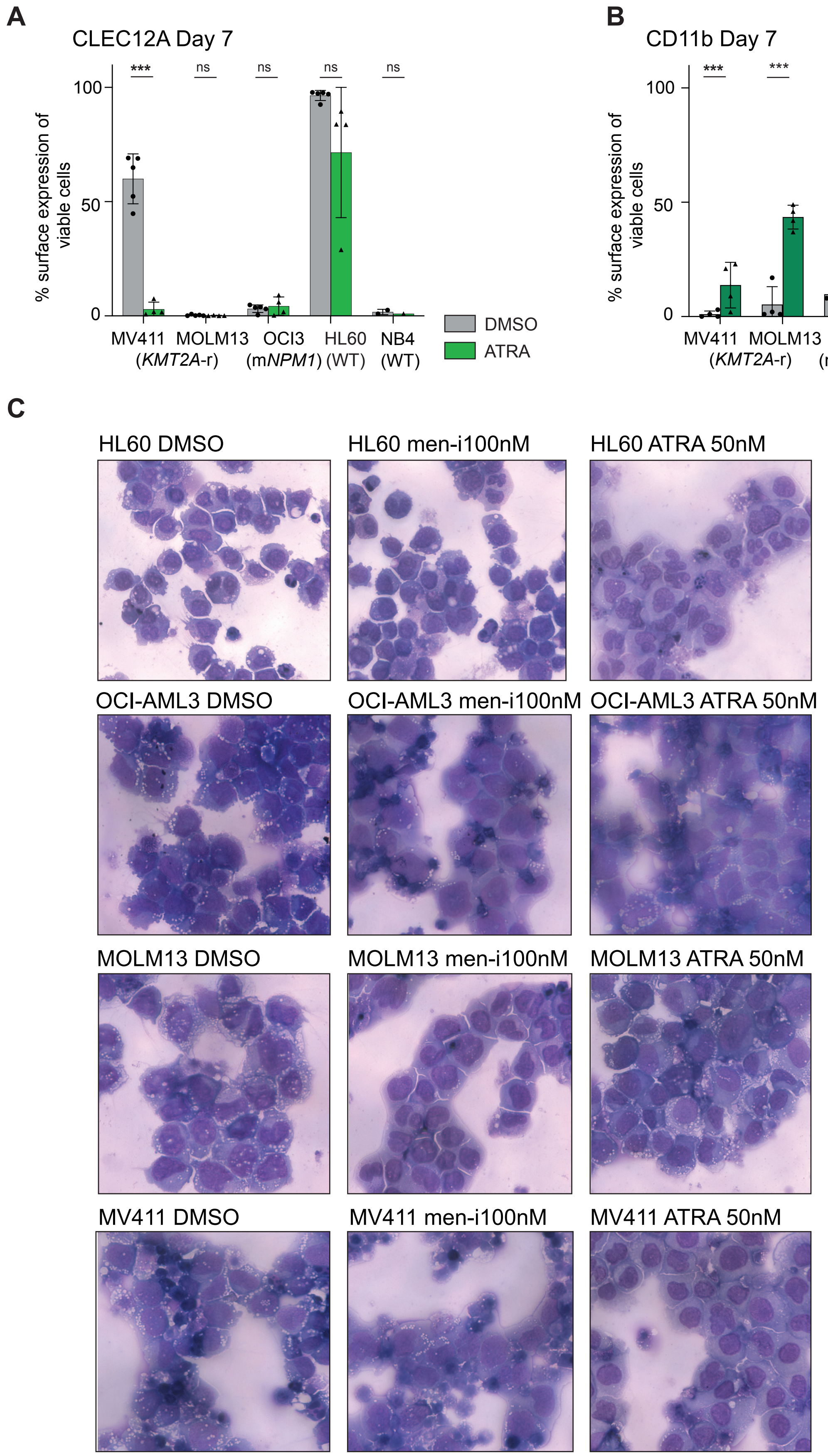

### Supplemental-Figure 3

**A** native T-cell proliferation stimulated with irradiated OCI-AML3

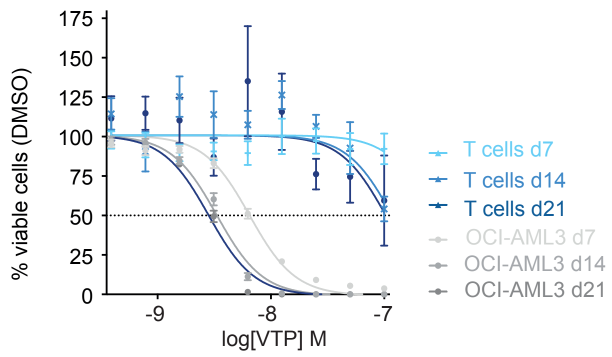


**B** Phenotype of native T-cells stimulated with irradiated OCI-AML3

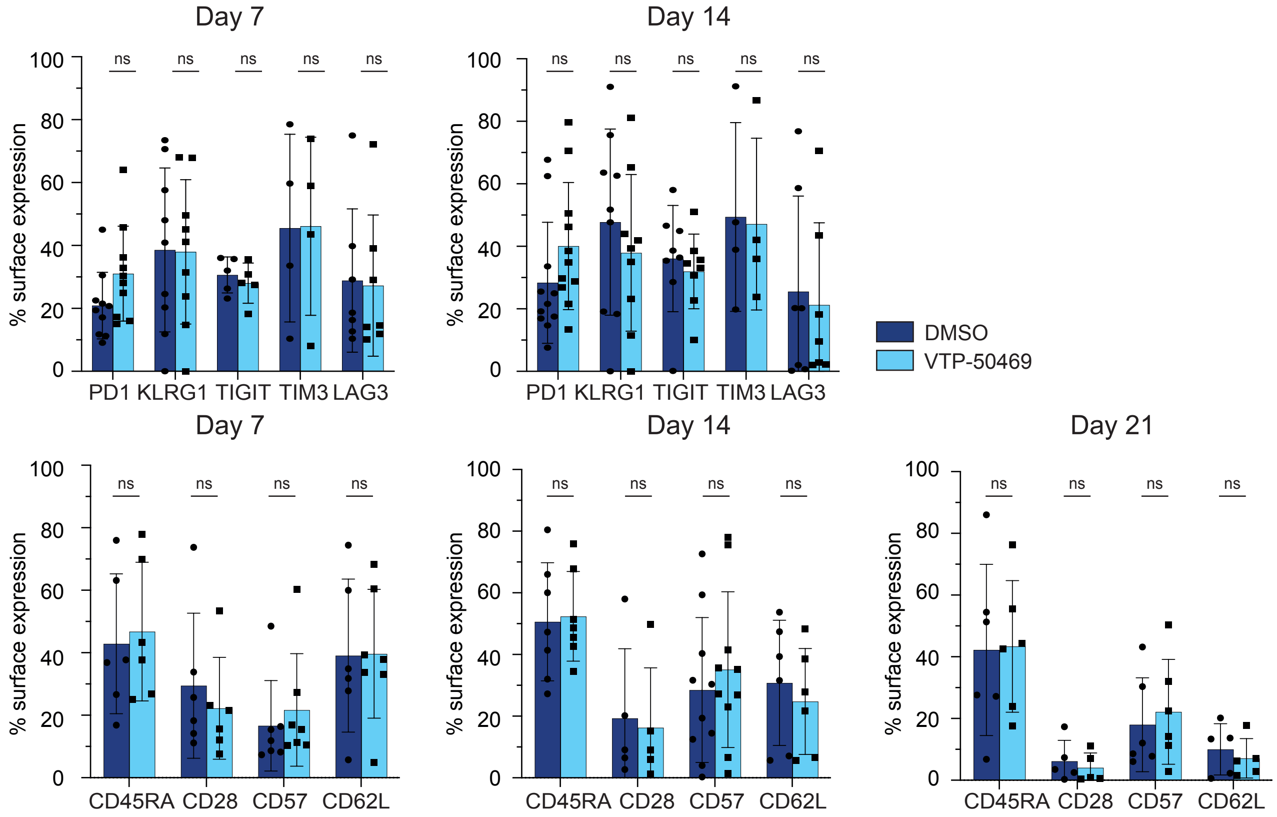

#### Supplemental-Figure 4

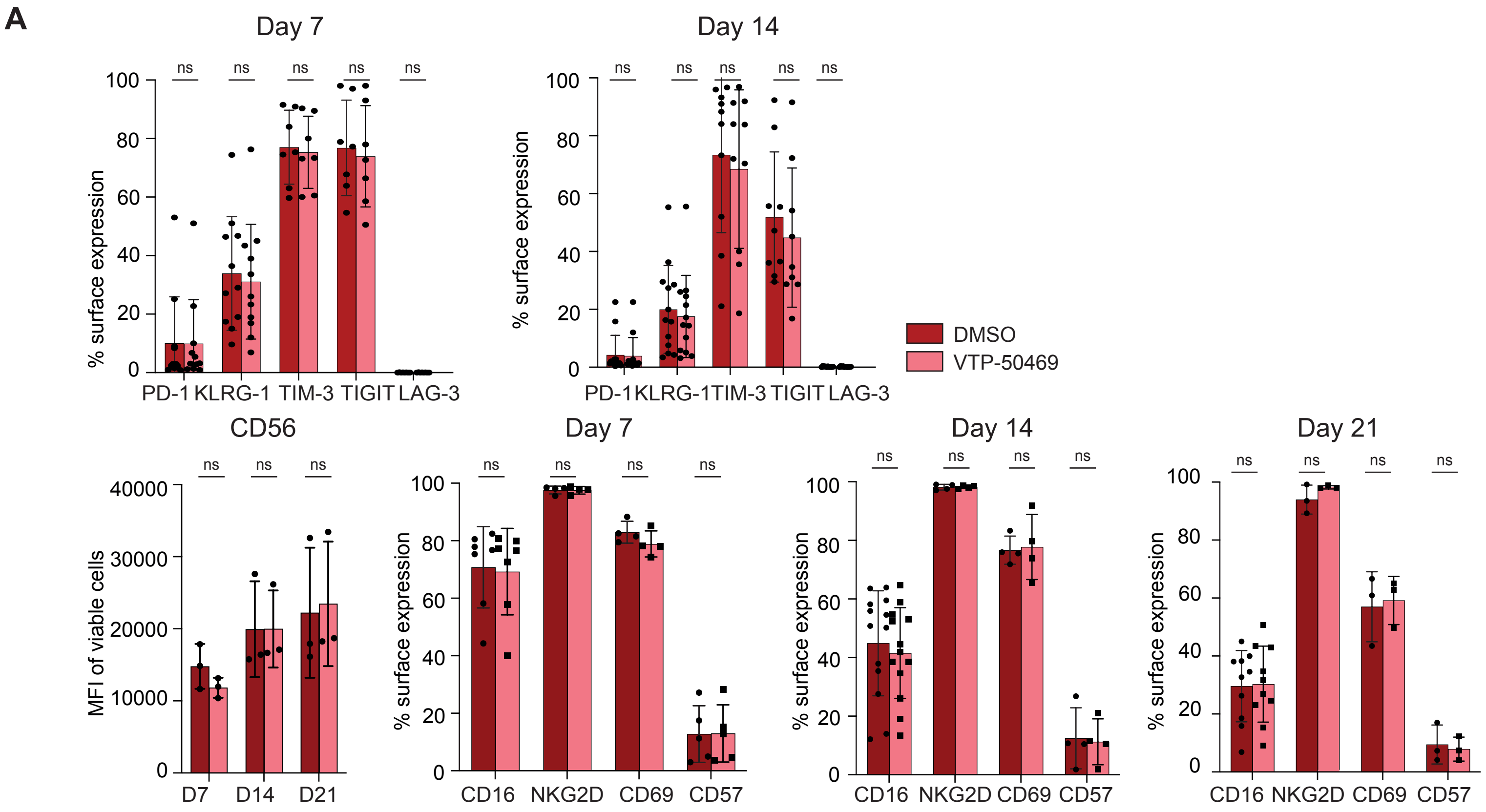

#### Supplemental-Figure 5

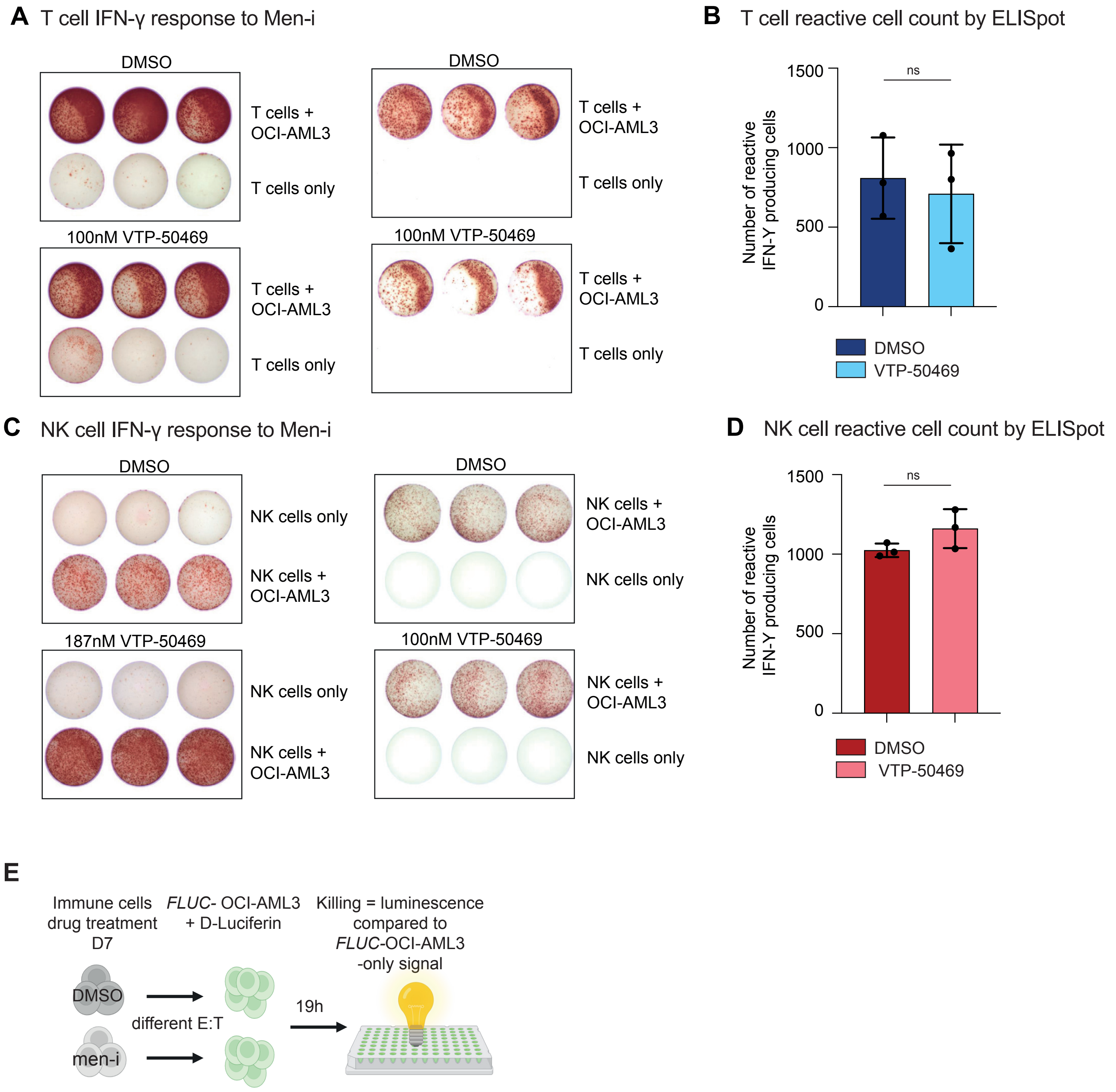

#### Supplemental-Figure 6

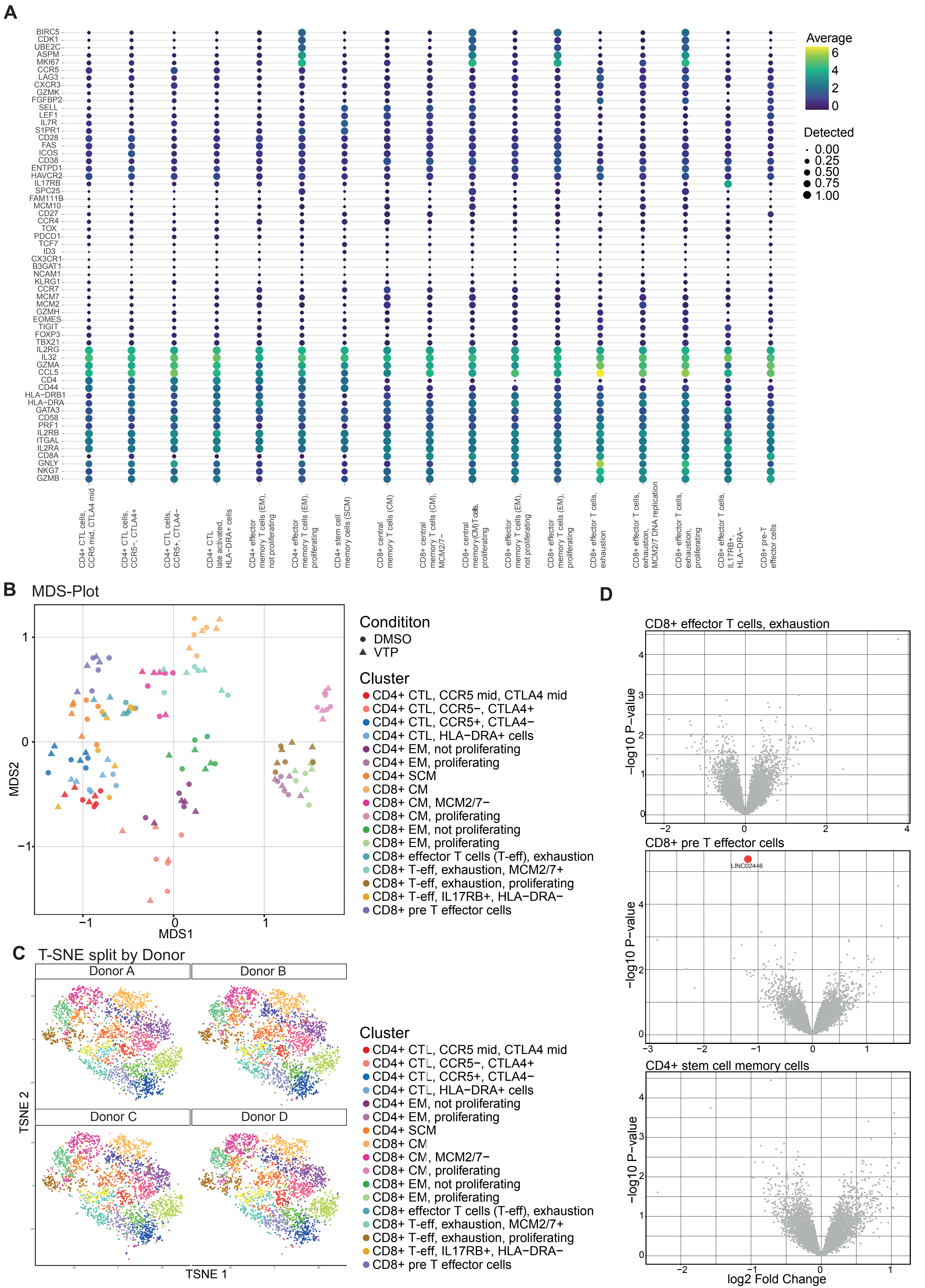

**Supplemental-Figure 7**

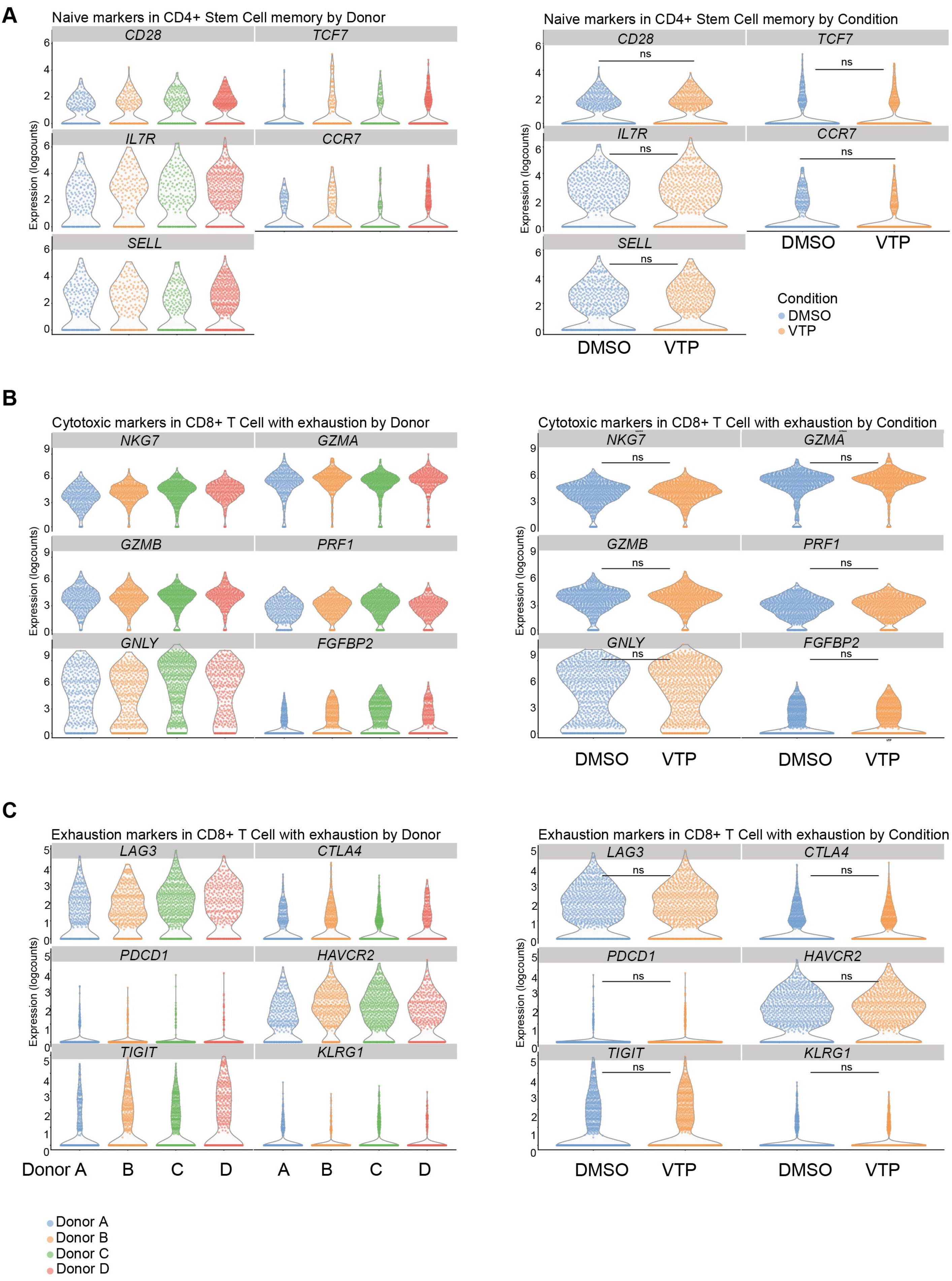

#### Supplemental-Figure 8

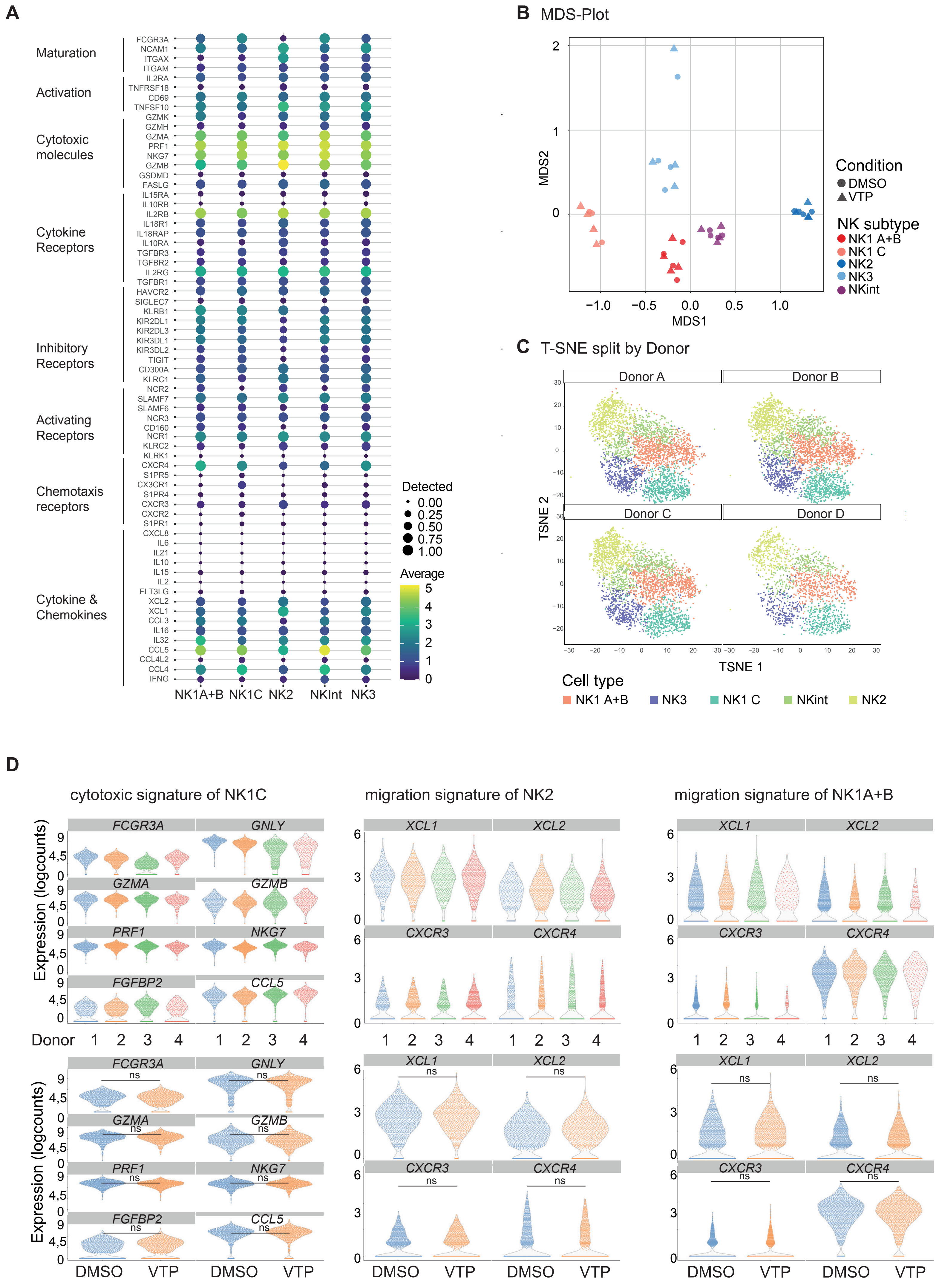

### Supplemental-Figure 9

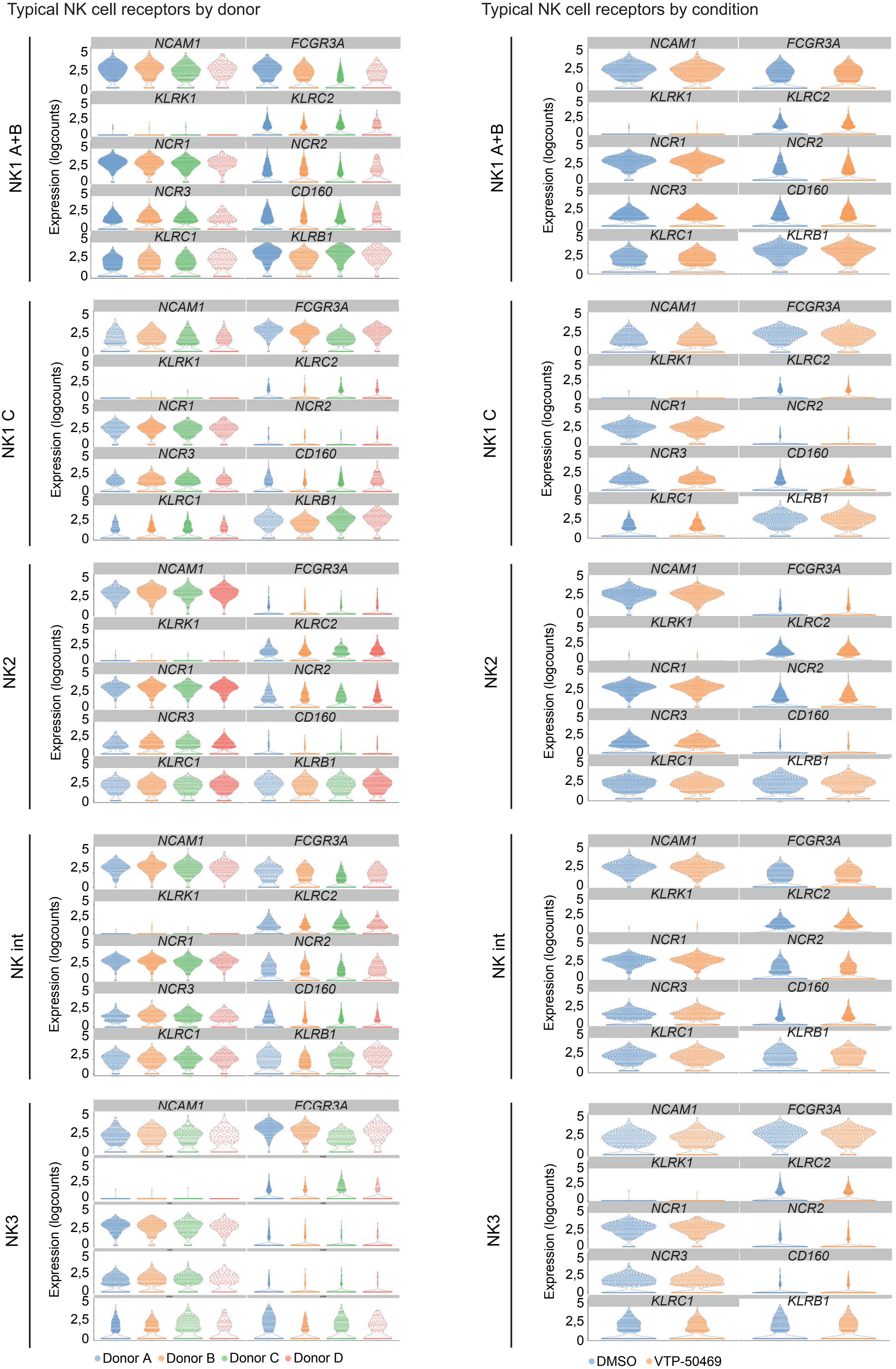

#### Supplemental-Figure 10

**A** Cytotoxicity of T cells against OCI-AML2 after 24h

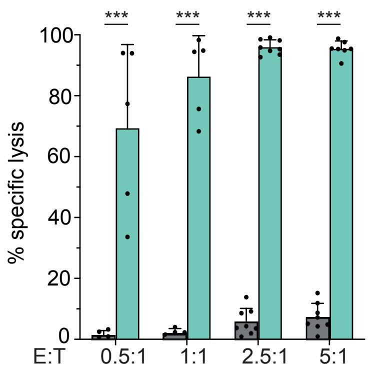


**B** Blast gating of primary cells for cytotoxicity assay with T cells

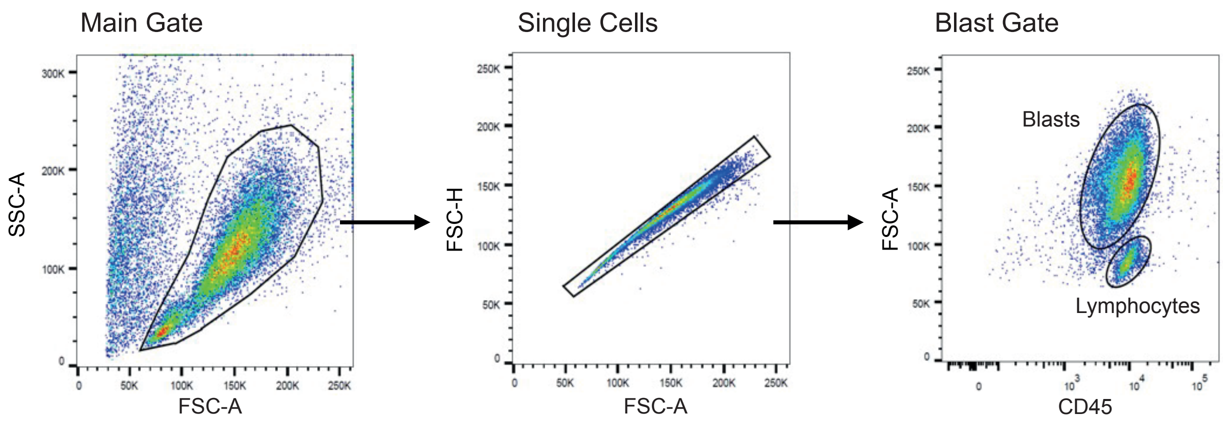


**C** CLEC12A expression of primary AML cells

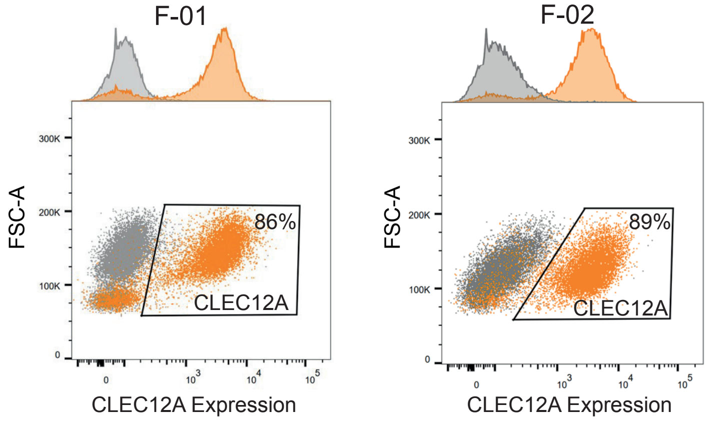

**Supplemental-Figure 11**

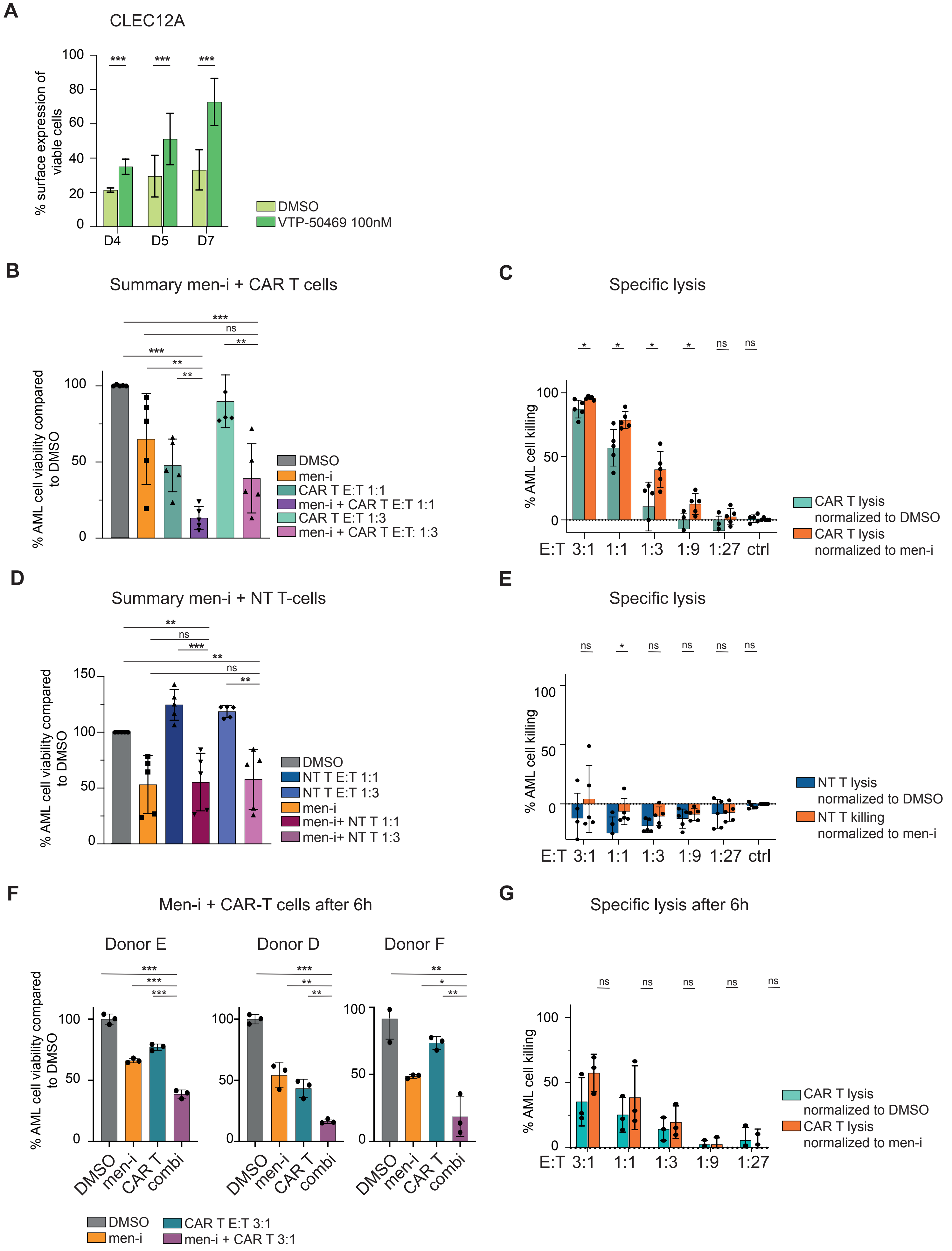

**Supplemental-Figure 12**

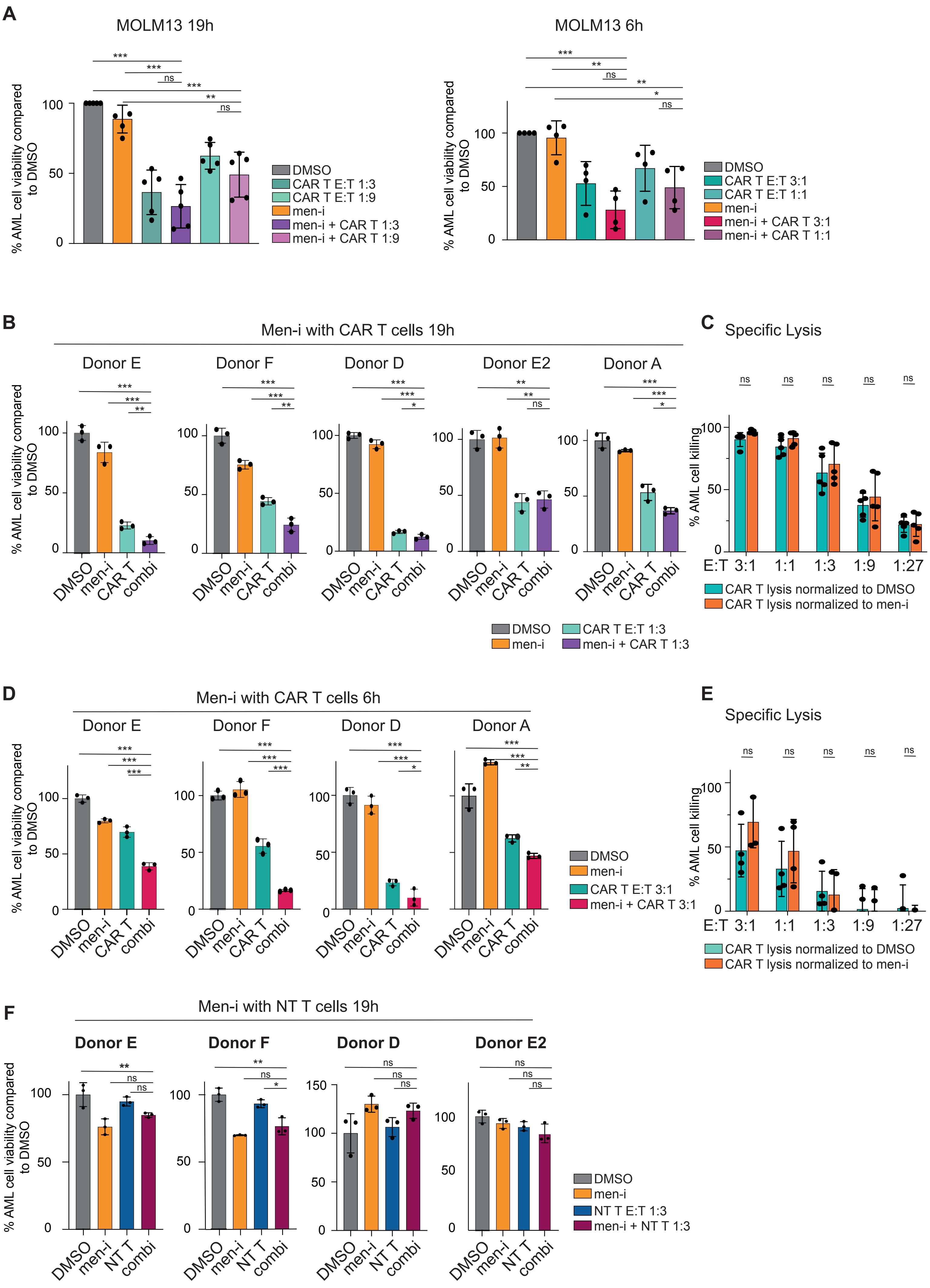

#### Supplemental-Figure 13

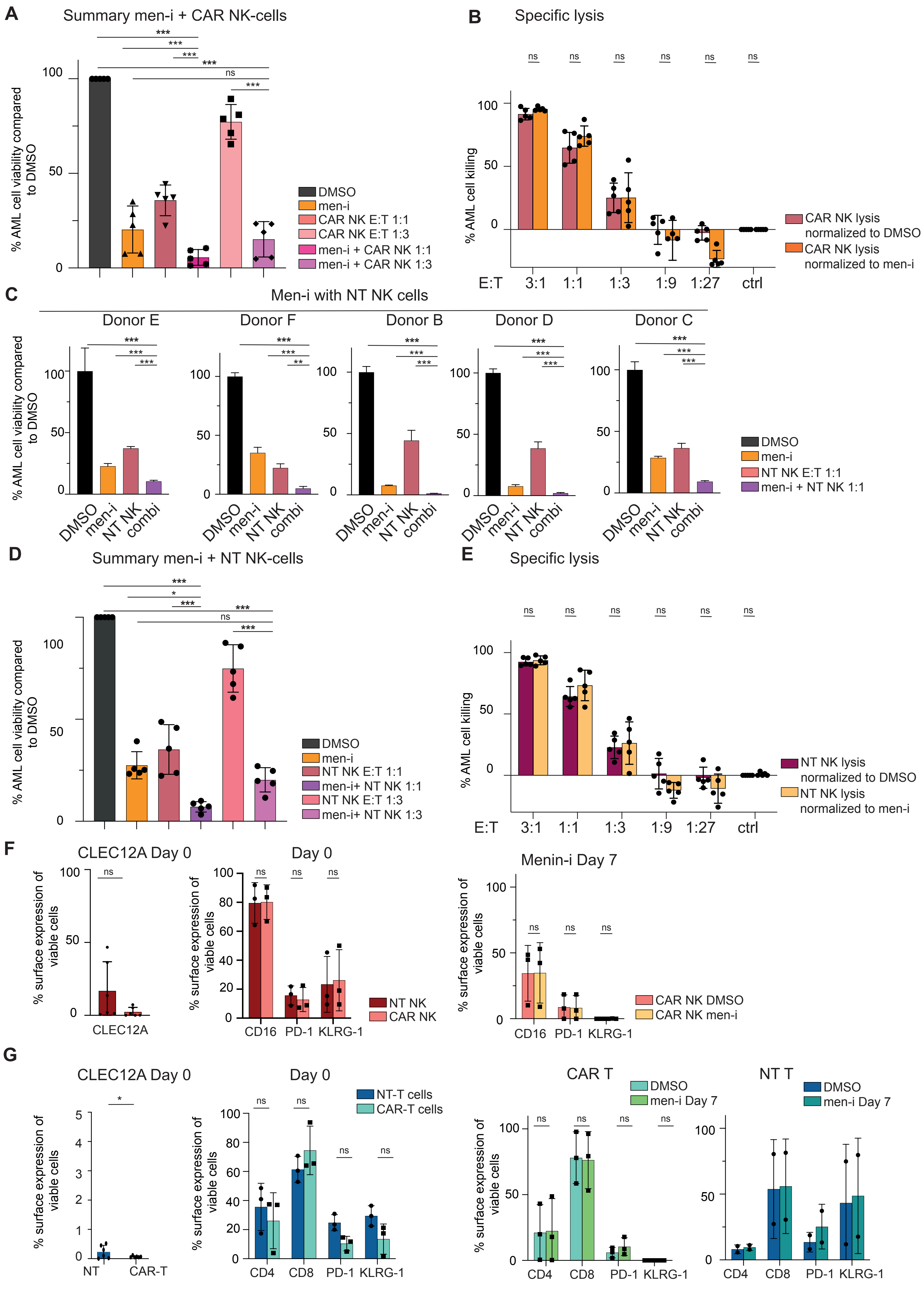

#### Supplemental-Figure 14

**A** *In vitro* efficacy of CAR T cell donor

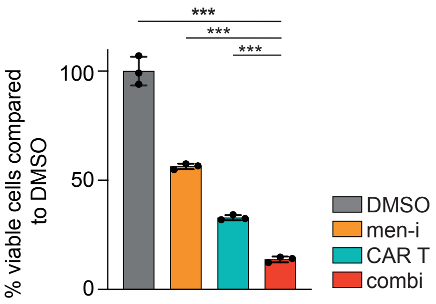


**B**

Gating of bone marrow engraftment (CD3 negative, CD45 positive)

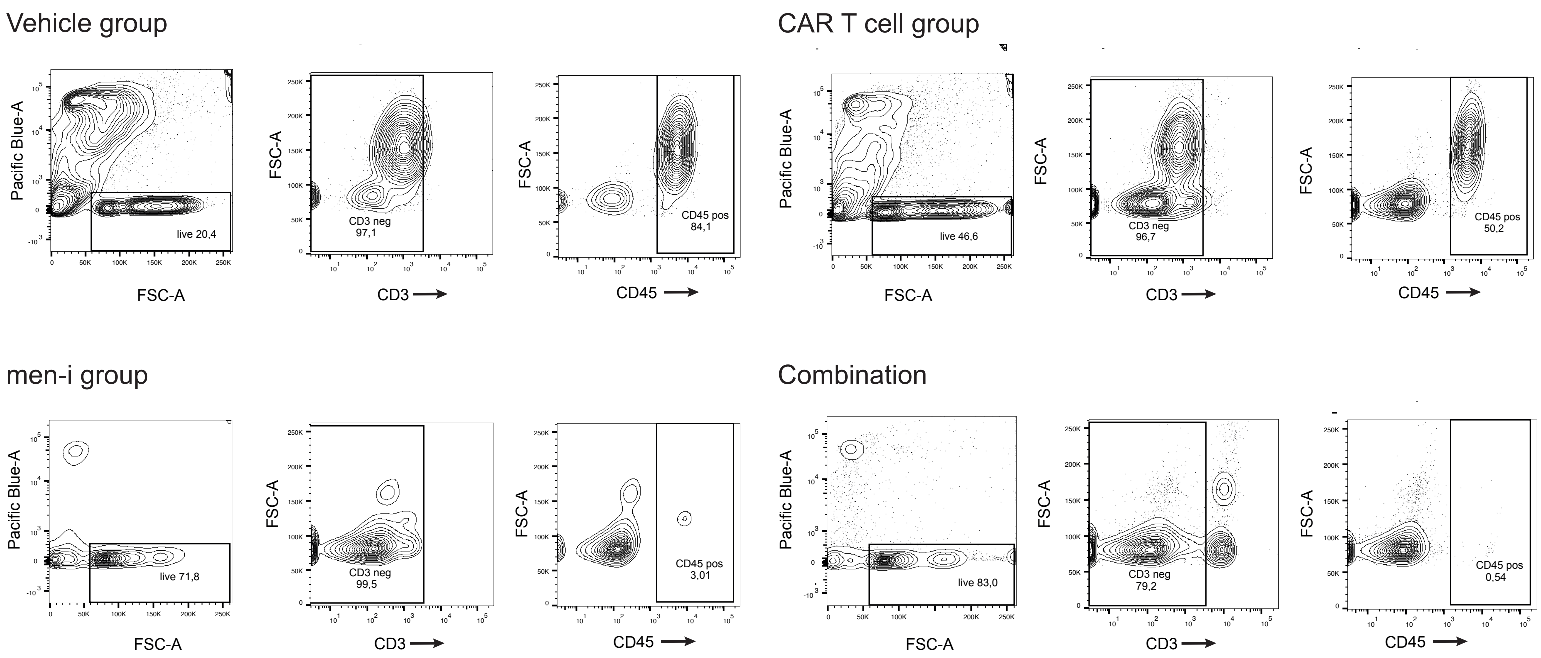


**C**

Engraftment day 10 (before CAR T cell injection)

Combination

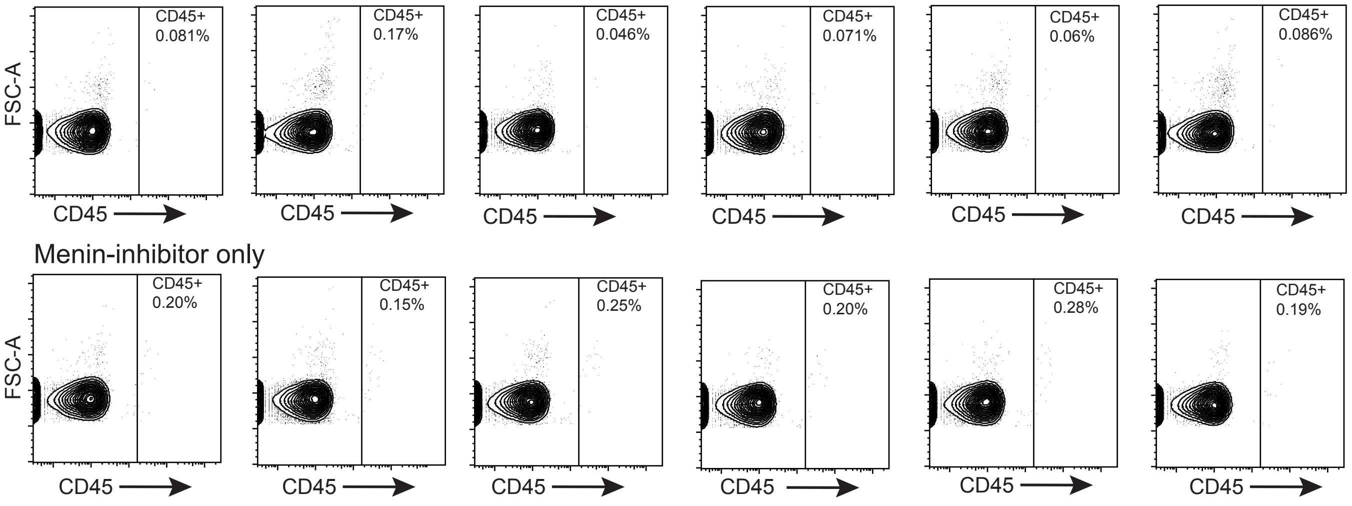
